# Obtusaquinone is a cysteine modifying compound that targets Keap1 for degradation

**DOI:** 10.1101/2020.04.03.023432

**Authors:** Christian E. Badr, Cintia Carla da Hora, Aleksandar B. Kirov, Elie Tabet, Romain Amante, Antoinette E. Nibbs, Evelyn Fitzsimons, John W. Chen, Norman Chiu, Ichiro Nakano, Wilhelm Haas, Ralph Mazitschek, Bakhos A. Tannous

## Abstract

We have previously identified the natural product Obtusaquinone (OBT) as a potent antineoplastic agent with promising in vivo activity in glioblastoma and breast cancer through the activation of oxidative stress; however, the molecular properties of this compound remained elusive. We used a multidisciplinary approach comprising medicinal chemistry, quantitative mass spectrometry-based proteomics, functional studies in cancer cells, pharmacokinetic analysis, as well as mouse xenograft models to develop and validate novel OBT analogs and charaterize the molecular mechanism of action of OBT. We here show that OBT and analogs, which have improved pharmacological properties, bind to cysteine residues with particular affinity to cysteine-rich Keap1, a member of the CUL3 ubiquitin ligase complex. This binding promotes an overall stress response and results in ubiquitination and proteasomal degradation of Keap1 and downstream activation of the Nrf2 pathway.

## INTRODUCTION

The transcription factor Nrf2 (Nuclear factor erythroid 2 (NFE2)-related factor 2) plays a key role in maintaining cellular homeostasis in response to oxidative stress by regulating the expression of antioxidant response element (ARE) dependent genes^1^. Keap1 (Kelch-like ECH-associated protein 1) is recognized as a predominant negative regulator of Nrf2, and functions as a substrate adaptor protein for the ubiquitin ligase CRL3 (cullin 3 (CUL3)-RING ubiquitin ligase). During homeostasis, Keap1 recruits Nrf2 to CUL3 thereby promoting its ubiquitination and subsequent proteasomal degradation. Nrf2 has been recognized to exert either pro-tumorigenic or anti-tumorigenic properties^2^. This apparent contradiction can be rationalized by differences in the cell state and the functional dependence on Nrf2 activation. Oncogenic activity is generally associated with constitutive activation of Nrf2 caused either by overexpression or somatic mutations of Nrf2 and/or other regulatory proteins^3^. Tumor suppressor activity, in contrast, is linked to transient activation of Nrf2, e.g. by small molecules^3, 4^. In this context, the Nrf2-Keap1 module has been validated and successfully pursued as a target for small molecules that disrupt Nrf2 binding and/or induce the dissociation of Keap1 from CUL3^5, 6^. These efforts have identified several inhibitor classes, including cysteine-reactive natural products and synthetic compounds, that bind Keap1 and activate the Nrf2 pathway ^7^.

We have previously identified the quinone methide Obtusaquinone (OBT), as a potent antineoplastic agent with selectivity over normal cells for glioblastoma (GBM), and several other cancer types^8^. Within the scope of this research, we have demonstrated that OBT treatment increases the production of reactive oxygen species (ROS) and induces DNA damage, leading to apoptotic cell death. However, the molecular mode of action of OBT, its medicinal chemistry, and *in vivo* pharmacology have not been understood, which has impeded preclinical development. We here developed novel OBT analogs with improved pharmacological properties and show that OBT is a thiol-reactive compound that reacts reversibly with cysteine residues, and particularly binds to Keap1, leading to CRL3-mediated autoubiquitination and proteasomal degradation of Keap1 thereby activating the Nrf2 pathway.

## RESULTS

### OBT forms reversible covalent adducts with thiols

OBT features a 2-hydroxy para-quinone methide, a moiety found in other natural products such as Celastrol^9^, which efficiently reacts with thiol nucleophiles including cysteine side chains to form substituted catechols (**Supplemental Fig. 1A**). It has previously been shown that the desmethoxy analog of OBT **SI1** (**Supplemental Fig. 1B**) reacts with glutathione (GSH) in aqueous buffer to form four distinct addition products, corresponding to the diastereomers derived from direct **SI2a,b** and vinylogous **SI3a,b** addition^10^. To demonstrate that OBT retains the ability to form sulfhydryl adducts, we incubated OBT with β-mercaptoethanol (BME) in ethanol and found that BME was readily added to OBT, preferentially (10:1) forming the direct addition product of the vinylogous addition product (**AF20**, **Fig. 1A**; **AF20a** and **AF20b**, **Supplemental Fig. 1C**). To investigate the reversibility of the thionucleophile addition to OBT, we incubated **AF20** in the presence of 5-fold excess cysteamine. Monitoring the reaction mix by LC/MS showed the formation of the corresponding amine functionalized analogs **SI4a** and **SI4b**, demonstrating that the addition of thiols is reversible or that the substitution proceeds through an S_N_2’ mechanism, and that the corresponding adducts exist in a dynamic equilibrium (**Supplemental Fig. 1C-D**).

**Figure 1:**
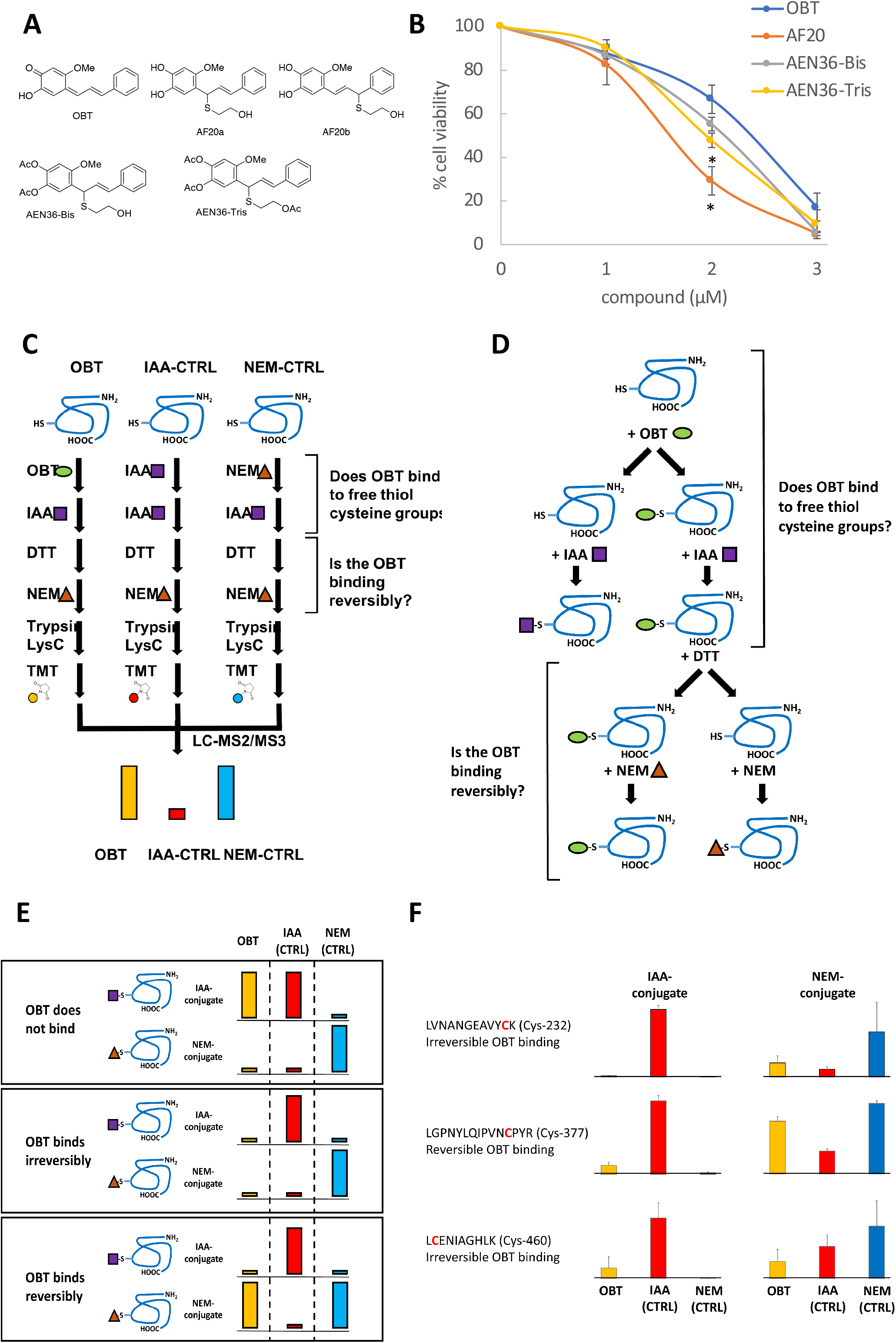
Characterization of OBT and its derivatives as reversible cysteine modifying drugs. **(A)** Chemical structure of OBT and novel analogs. **(B)** MDA-MB231 cells were treated with different doses of OBT or analogs in 4-replicates and cell viability was measured four days later and expressed as % of vehicle control. *P<0.05 Student *t* test versus OBT. **(C)** An overview of the mass spectrometry-based approach to interrogate OBT binding to cysteine residues. OBT is given to the protein and will or will not react free thiol groups. In a second step, an alkylating reagent, iodoacetamide (IAA), is added to the mixture, which will react with free thiols if they are not blocked by OBT. In a third step a reductant, dithiothreitol (DTT), is added to the mixture. If OBT binding is reversible under these reductive conditions, free thiol groups will be available for reaction with another alkylating reagent, N-ethylmaleimide (NEM), added in the last step. The combined reaction product is digested, and peptides are subjected to mass spectrometry followed by multiplexed quantitative proteomics to determine for each cysteine if OBT did bind to it and if this binding can be reversed; results are compared to 2 controls (1) sample with protein alkylated only with IAA and (2) sample alkylated only with NEM. Each cysteine-containing peptide will reveal a certain intensity pattern across the three samples that will allow to determine site-specific binding activity of OBT. (**D**) Potential outcomes for OBT treated sample in the differential alkylation reaction process. (**E**) Multiplexed mass spectrometry is used to quantify peptides containing only cysteine residue generated from a consecutive trypsin and LysC digest across all samples. Quantifying IAA and NEM modified peptides across all three samples produces unique patterns of peptide intensities for each of the three options: (i) OBT does not bind, (ii) OBT binds irreversibly, and (iii) OBT binds reversibly. (**F**) Differential alkylation was used on bovine catalase to study OBT-binding properties. The IAA conjugate intensity patterns showed that OBT reacted with all three identified cysteines (Cys-232, 377, and 460). The NEM-conjugate patterns showed that binding was irreversible for cysteines 232 and 460 but reversible for cysteine 377 under these experimental conditions.

Next, we explored if the reactivity with thiols could be exploited for the development of novel OBT analogs with improved pharmacological properties to overcome the limited solubility of OBT in aqueous media, and to buffer the abrupt oxidative stress as a result of rapid depletion of the intracellular GSH pool caused by thiol-reactive compounds ^11^. We have previously shown that addition of GSH or N-acetyl cysteine (NAC) reduces the activity of OBT in cell culture, likely by extracellular scavenging of OBT^8^. Based on our hypothesis that the ability to react with cysteine side chains is critical for OBTs activity, we postulated that prodrugs designed to liberate OBT or analogs that retain the ability to react with cysteines would yield improved inhibitors, including compounds with enhanced solubility, while mitigating the general toxicity as a result of GSH depletion. AF20, the adduct of OBT and BME, which is more soluble and liberates OBT in PBS, potently killed patient-derived glioma stem-like cells (GSCs) neurospheres, while exhibiting lower toxicity towards primary human astrocytes (HA) (**Supplemental Fig. 2A**)^8^. As previously observed with OBT, the activity of AF20 is reversed in the presence of NAC (**Supplemental Fig. 2B**). These results are consistent with our proposed mechanism that AF20 converts to OBT as active compound and that NAC may not only function as a ROS antagonist but also as a direct scavenger of OBT.

To block the direct reversibility observed with AF20, we speculated that acetylation of the catechol would stabilize the thiol adduct in aqueous media but predicted that the phenolic esters would be cleaved intracellularly allowing for intracellular release of OBT via AF20 or following displacement by an S_N_2’ mechanism (See supporting information). Treatment of AF20 with 2eq and 3eq acetic anhydride yielded AEN36 Bis and AEN36 Tris (**Fig. 1A**), respectively. Both compounds demonstrated increased stability and increased potency on different cancer cells as compared to OBT (**Fig. 1B** **and** **Supplemental Fig. 2C**).

To confirm reversible cysteine modification by OBT, we established a mass spectrometry-based approach that would allow the identification of specific cysteine side chains in native proteins that covalently react with OBT and differentiate reversible from irreversible interactions. The experimental set-up is outlined in **Fig. 1C-D**, and the potential outcome **in Fig. 1E**. The method is based on serial exposure of a protein to OBT, the cysteine-alkylating reagents iodoacetamide (IAA) to monitor derivatization with OBT, to dithiothreitol (DTT) to probe reversibility of OBT binding, and to another alkylating reagent, *N*-methylmaleimide (NEM), to allow a readout of probing the reversibility. In two parallel reactions, OBT is replaced by IAA and NEM. These reactions are used as standards to enable a final reaction read-out by multiplexed quantitative mass spectrometry^12, 13^. We performed this experiment using bovine catalase, a cysteine-rich antioxidant (**Supplemental Fig. 2D)**, and found that OBT binds to all detected peptides with cysteine residues (**Fig. 1F**; **Supplemental Tables 1-2**). For Cys-377, where the cysteine is followed by a proline in the protein sequence, we observed reversible binding, while the two other monitored cysteine residues (Cys-232 and Cys-460), showed tight or irreversible binding under the tested reaction conditions (**Fig. 1F**). These results show that OBT is an effective cysteine-alkylating reagent and that the alkylation is reversible but might depend on structure of peptide or amino acids adjacent to cysteine residue, binding affinity and/or dissociation kinetics. It is to be noted that reversibility was determined under the condition of 2-fold access of DTT over alkylating reagents and that OBT binding may be affected differentially under other conditions.

### OBT activates the Nrf2 pathway *in vitro* and *in vivo*

To gain a better insight into the molecular mechanism of OBT and its effect on the global proteome, we implemented a multiplexed quantitative mass spectrometry-based proteomics using the isobaric labeling strategy with Tandem Mass Tag (TMT) reagents and the MS3 method^12, 14^. Global proteomics analysis in two different patient-derived GSCs specimens at 20h after treatment with a subtoxic dose of OBT, identified heme oxygenase 1 (HMOX1; HO1) as the top upregulated protein in both lines (**Fig. 2A-B**; **Supplemental table 3**). Gene Ontology category analysis of the most upregulated proteins after treatment (using DAVID bioinformatics platform) showed that the most significantly enriched category was “oxidation-reduction process” containing the upregulated proteins HMOX1, PHS2, GLRX1, VKORL, BLVRB, and CDO1. To further explore the functional network of these proteins, we interrogated the STRING database for high-confidence direct interactors of these proteins (white circles; **Fig. 2C**), and found a network of densely connected proteins containing five of the upregulated proteins (red) as well as Nrf2 (green), a master regulator of oxidative damage response and Keap1 (green), the E3-ligase regulating the ubiquitin-mediated degradation of Nrf2 (**Fig. 2C**). A strong increase in HO1 mRNA expression in response to OBT was detected across different cancer cells (**Fig. 2D** **and** **Supplemental Fig. 3A**). The Nrf2 target genes NQO1 and TXNRD2 were also upregulated following OBT treatment (**Supplemental Fig. 3B**). To further support these findings independently, we designed a functional ARE luciferase reporter which was strongly activated following treatment with sub-toxic doses of OBT, even higher than the positive control *tert*-butylhydroquinone (tBHQ), a potent activator of Nrf2^15^ (**Supplemental Fig. 3C-D)**. Importantly, OBT-induced ARE activation and loss of cell viability was completely reversed by the addition of various thiol nucleophiles antioxidants including NAC, dithiothreitol (DTT), GSH, but only partially by Trolox, an antioxidant that is devoid of thiol groups (**Fig. 2E** **and** **Supplemental Fig. 3E**). Finally, we confirmed OBT-mediated Nrf2 activation *in vivo* using a breast cancer mouse model generated by injecting MDA-MB-231 into the fat pad of nude mice^8^. Treatment with OBT (7.5mg/kg for four consecutive days) resulted in 7-fold upregulation of HO1 transcripts in the tumor, consistent with our findings in cell culture (**Fig. 2F**). Taken together, these results confirm that OBT acts as an inducer of Nrf2 both in culture and *in vivo*.

**Figure 2:**
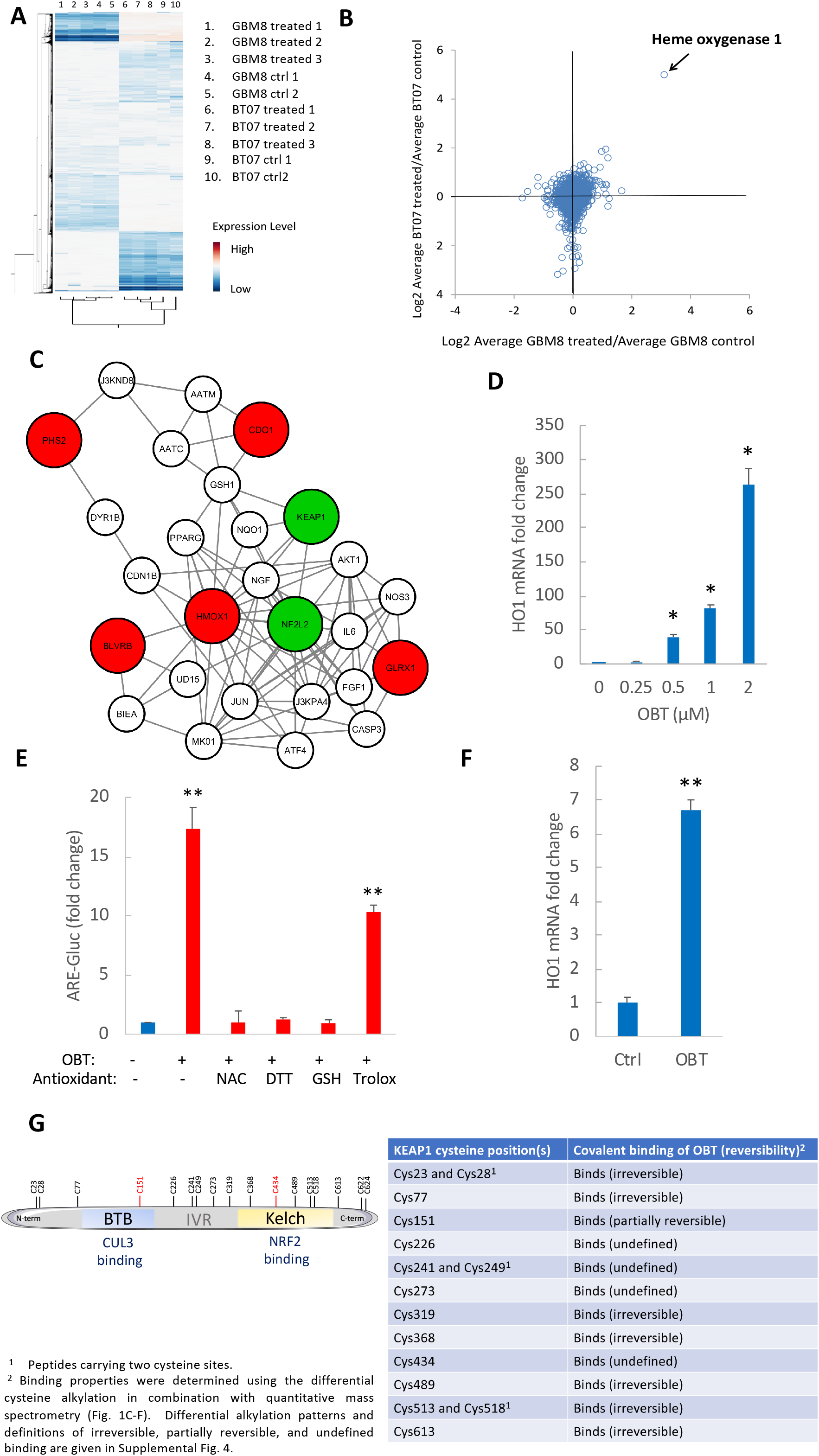
OBT activates Nrf2 pathway *in vitro* and *in vivo*. (**A-C)** Quantitative mass spectrometry-based proteomics post-OBT treatment using 10-plexed tandem mass tags (TMT) to simultaneously map protein concentration changes of 7,904 proteins in two GSCs, GBM8 and BT07, either treated with OBT (triplicates) or untreated (duplicates) for 20 hours. **(A)** heat map derived from unsupervised hierarchical clustering of the data. **(B)** Relative protein concentration differences between OBT treated and control cells. **(C)** Functional network of upregulated proteins following OBT treatment using STRING database showing high-confidence direct interactors with HMOX1 (white circles), protein connection containing five of proteins upregulated in the OBT-treated samples (red) as well as Nrf2 (green). **(D)** HO1 mRNA levels were determined by qRT-PCR (normalized to GAPDH) in MDA-MB-231 cells treated with the indicated doses of OBT for 8 hours. **(E)** U87 cells stably expressing *Gaussia* luciferase under the control of ARE response elements (ARE-Gluc) and the constitutively active *Vargula* luciferase (Vluc under control of SV40 promoter for normalization of cell number) were treated with OBT and different antioxidants. Sixteen hrs post-treatment, aliquots of the conditioned medium were assayed for Gluc and Vluc activities and data were expressed as ratio of Gluc/Vluc, normalized to the control (set at 1). **(F)** Mice carrying fat pad MDA-MB231 tumors were treated with a daily dose of either DMSO or OBT (7.5mg/kg, intraperitoneally) for four consecutive days. At day 5, tumors were removed, RNA was extracted and analyzed for HO1 mRNA by qRT-PCR (normalized to GAPDH). **P<0.05 Student *t* test versus control. **(G)** Mapping of OBT covalent biding to cysteine residues in Keap1. OBT was found to bind covalently to all identified cysteine sites and this modification was found to be partially reversible at two sites (Cys151 and Cys434).

### OBT is a cysteine-modifying drug targeting Keap1

Several thiol reactive Nrf2 activators have been shown to bind Keap1, suggesting that OBT could function in similar fashion ^16^. To investigate if Keap1 is a target of OBT, we applied the mass spectrometry-based approach described in Figure 1 to Keap1. Similar to the experiment with bovine catalase, we found OBT to bind covalently to all identified cysteine sites, thus confirming that this compound can directly bind to Keap1. Importantly, we found this modification to be partially reversible at Cys151 located in the BTB domain (CUL3 binding site) and Cys434 located in the Kelch domain (Nrf2 binding site) of KEAP1 (**Fig. 2G**; **Supplemental Figure 4** **and Table 4-5**).

Next, we evaluated the effect of OBT treatment on cells following stable downregulation of Keap1. Silencing of Keap1 with shRNA (shKeap1) expectedly resulted in stabilization of Nrf2, leading to a major increase in ARE reporter activity (**Fig. 3A-B** **and** **Supplemental Fig. 5A**). There was no strong potentiation of OBT-mediated ARE activation in cells expressing shKeap1 as compared to those expressing a non-targeting shRNA (shCtrl), detected using the ARE reporter (**Fig. 3A** **and** **Supplemental Fig. 5B**) and mRNA expression of HO1 and NQO1 (**Supplemental Fig. 5C)**. Further, silencing of Keap1 decreased cell death following treatment with OBT (**Fig. 3C** **and** **Supplemental Fig. 5D**). These data suggests that either Keap1 expression is necessary for binding of OBT to Keap1 cysteine residues, thus stabilizing and activating Nrf2, or that a strong activation of Nrf2 protects against OBT-mediated cytotoxicity.

**Figure 3:**
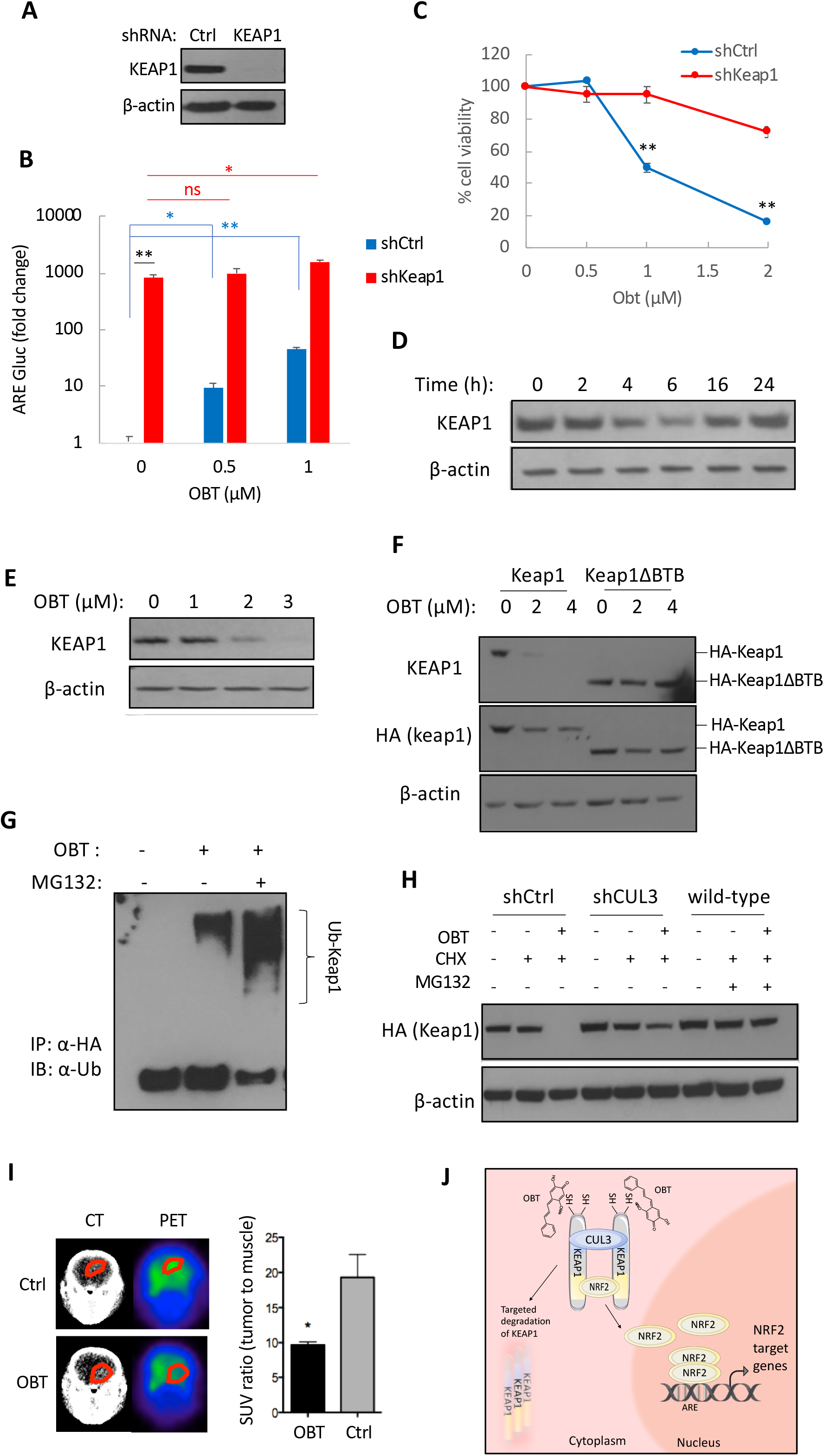
OBT targets Keap1 and induces its degradation. **(A)** Western blot analysis of Keap1 protein in U87 cells stably expressing shCtrl or shKeap1. **(B)** ARE-Gluc activity (normalized to Vluc) sixteen hrs post-OBT treatment in U87 glioma cells expressing a non-targeting shRNA (control; shCtrl) or shKeap1. (**C**) U87 cells stably expressing shCtrl or shKeap1 were treated with different doses of OBT and cell viability was measured after four days. (**D**) U87 cells were treated with 1 μM OBT and cell lysates were collected at the indicated time points and analyzed for Keap1 protein levels by Western blotting with β-actin as loading control. **(E)** Keap1 protein levels in U87 cells treated with the indicated doses of OBT for 8 hours. **(F)** 293T cells transfected with a plasmid expressing HA-tagged Keap1 or Keap1ΔBTB were treated with OBT (2 or 4 μM) or control. Cell lysates were collected after 6 hours and immunoblotting was performed using anti-Keap1 and anti-HA antibodies. **(G)** 293T cells were treated with OBT (2 μM) and/or MG132 (10 μM) for 6 hours before immunoprecipitation with anti-HA antibody. Ubiquitination was determined using anti-Ubiquitin antibody. **(H)** MDA-MB-231 cells (wild-type) or same cells stably expressing shCtrl or shCUL3 were transfected with HA-Keap1 and treated with OBT (3 μM), CHX (3 μg/mL) and/or MG132 for 6 hours. Keap1 levels were detected using anti-HA. **(I)** Mice-bearing patient-derived GBM8 GSC tumors were treated with 3 doses of OBT (10mg/kg within a 24h period) or vehicle control (n=5/group) and imaged 4h after the last OBT administration. Representative FDG-PET-CT scans (left) and signal quantification (right) showing that OBT lead to a 50% decrease in FDG tumor uptake, determined using tumor to muscle ratio with muscle as the background. \**P*<0.05; **P<0.001 Student *t*-test. **(J)** Schematic representation of the proposed mode of action of OBT-mediated degradation of Keap1, resulting in subsequent nuclear translocation of Nrf2 to bind ARE and activate transcription of downstream targets.

Numerous anticancer drugs have been shown to activate Nrf2^16^. We tested if reactive electrophiles and oxidants known as transient activators of Nrf2 could induce cell death when added to breast and brain cancer cells, similar to OBT. We first confirmed ARE-inducing properties of the triterpenoid CDDO-Me (2-cyano-3,12-dioxoolean-1,9-dien-28-oic acid-methyl ester)^17^, currently being clinically tested for the treatment of leukemia and solid tumors as well as other diseases. U87 and MDA-MB-231 cells expressing ARE-Gluc reporter treated with CDDO-Me showed increased reporter activity **(Supplemental Fig. 6A)**. Higher doses of this compound led to a marked decrease in cell viability in U87 cells **(Supplemental Fig. 6B)**. Treatment of U87 cells with additional Nrf2 activators, Cinnamaldehyde^18^, diethyl fumarate^19^ and sulforaphane^20^ also resulted in increased ARE activity and a moderate decrease in U87 cell viability at the doses tested **(Supplemental Fig. 6C-D)**. When combined with OBT, all three compounds showed increased cytotoxicity **(Supplemental Fig. 6E)**. These results suggest that increased electrophile or oxidant concentrations is likely to cause further cysteine modifications in the cellular proteome along with an increase stress response as evident by an upregulation of Nrf2 signaling, thus increasing cytotoxicity.

### OBT promotes the degradation of Keap1

We next asked whether OBT-mediated covalent modification of Keap1 affects stability of this protein. Immunoblot analysis showed a time- and dose-dependent decrease in Keap1 protein levels as early as 4h following treatment with OBT, which were restored to physiological levels at 24 h (**Fig. 3D-E**). To further confirm Keap1 protein degradation, we expressed HA-tagged Keap1 or Keap1 with a deleted BTB domain (keap1ΔBTB)^21^ which is essential for Keap1 binding to CUL3 and activation of Keap1-CUL3 E3 ligase activity. We did not observe any decrease in Keap1 protein levels in cells expressing BTB-mutated Keap1 (**Fig. 3F**), suggesting that the E3 ligase activity is required for OBT-mediated degradation of Keap1. Further, similar to Keap1 knockdown, silencing of CUL3 with shRNA prevented ARE activation following treatment with OBT (**Supplemental Fig. 5B**) and decreased cell death (**Supplemental Fig. 5D**). Ectopic expression of a dominant-negative CUL3 mutant (DN-CUL3) also protected against OBT-induced cell death, further corroborating these findings (**Supplemental Fig. 7A**). The neddylation of CUL3 is essential for its ubiquitin ligase activity^22^. To determine if CUL3 activation is essential for OBT-induced cell death, we co-treated cells with OBT and the neddylation inhibitor MLN4924 and observed protection against cell death (**Supplemental Fig. 7B**). Overall, these findings confirm that E3 ligase activity is required for targeting of tumor cells with OBT.

Since ubiquitination of Keap1 could lead to its degradation^23^, we evaluated this process following OBT treatment by immunoblotting. Indeed, OBT treatment effectively resulted in ubiquitination of Keap1 (**Fig. 3G**). Additionally, downregulation of CUL3 or co-treatment with the proteasome inhibitor MG132 prevented OBT-mediated degradation of Keap1 (**Fig. 3H**). Among the reactive cysteines of Keap1, C151 was found to be necessary for Keap1-alkylating ARE inducers, which, promote the dissociation of the Keap1-CUL3 complex, thus stabilizing Nrf2. Accordingly, mutation of C151 impairs its alkylation by electrophiles such as sulforaphane, tBHQ or AI-1 and impairs Nrf2 activation^24–26^. However, serine substitution of Cys-151 (Keap1C151S) did not protect against OBT-mediated degradation of Keap1 (**Supplemental Fig. 7C**). Overall, these results confirm that OBT treatment leads to degradation of Keap1 and that CUL3 is an essential regulator of this process.

### OBT and its analog effectively target tumors in preclinical mouse models

Pharmacokinetic profiling of OBT in mice showed high systemic plasma clearance with terminal plasma half-life of 24 minutes with intraperitoneal injection (**Supplemental Fig. 8A**). Furthermore, we found that OBT efficiently penetrates the intact blood-brain barrier (BBB) (**Supplemental Fig. 8B-C**). To confirm that OBT is able to penetrate the brain and functionally target brain tumors, we used positron emission tomography (PET) with the PET-tracer 2 [^18^F] fluoro-2-deoxy-D-glucose (FDG), which measures the rate of tumor glucose uptake in a mouse orthotopic GSCs model. OBT-treated mice exhibited approximately a 50% decrease in FDG tumor uptake as measured by FDG-PET imaging (**Fig. 3I**). Finally, we evaluated the *in vivo* antineoplastic effect of one of the analogs, AEN36 Tris, in a breast cancer mammary fat pad tumor xenograft model. Treatment with this OBT analog (10mg/kg daily for 22 days) induced a significant decrease in tumor volume, compared to the control group, as assessed by bioluminescence imaging (**Supplemental Fig. 8D)**. In summary, OBT penetrates the BBB and can be modified to enhance its solubility while retaining its antineoplastic properties.

## DISCUSSION

Small molecules that react with cysteine side chains within Keap1^26^ or target the kelch domain of Keap1^27^ have been identified. We now provide direct evidence that the natural compound OBT activates the Nrf2 pathway by binding covalently to cysteine residues within the BTB-domain, of Keap1 leading to its ubiquitination and subsequent proteasomal degradation. This directly impacts the ability of the CUL3-Keap1 ubiquitin ligase complex to degrade Nrf2, resulting in Nrf2 stabilization and downstream activation of ARE-mediated transcription (**Fig. 3J**). It is highly likely that OBT also interacts with other thiol-rich proteins; however, our data supports that Keap1 is a major functional target for OBT and that the BTB-CUL3 ubiquitin ligase complex is required for OBT-mediated degradation of keap1. It is reasonable to assume that this cysteine-reactive compound likely engages other secondary targets in order to promote an overall stress response. In fact, downregulation of Keap1 was not sufficient to induce the same level of cytotoxicity observed after treatment with OBT or other Nrf2 activators.

The transcription factor Nrf2 is often viewed as a pleiotropic gene. Whether its activation or inhibition is beneficial for tumor treatment remains a paradox and seems to depend on various factors such as the cell type, tumor stage, and genetic aberrations within the tumor ^2–4^. Nrf2 has been suggested to act as a tumor suppressor, thus its activation can suppress carcinogenesis^2, 28–30^. On the other hand, Nrf2 activation was shown to decrease tumor growth in established tumors^31^, and several antineoplastic drugs enhance Nrf2 activity^16^. In this study, we have demonstrated that, in addition to OBT, several other reactive electrophiles and oxidants known as transient activators of Nrf2 can induce cytotoxicity in tumor cells. On the other hand, activation of Nrf2 by cancer targeting drugs can also lead to unfavorable clinical outcomes because of Nrf2 ability to enhance chemoresistance^28^. One plausible explanation of this paradox is that, unlike somatic mutations and oncogene-mediated signaling that promote a sustained Nrf2 activation accompanied by numerous adaptation mechanisms, pharmacological activation of Nrf2 is transient and does not neceserrily phenocopy constitutive Nrf2 activation^3, 4^. Despite these controversies, several Nrf2 activators have been developped as antineoplastic compounds in preclinical studies^32^, and at least one such compound, sulforaphane, has advanced to a phase 2 clinical trial for the treatment of metastatic breast cancer (NCT02970682).

The dose-response curve of many chemopreventive agents as well as chemotherapeutic drugs is U-shaped^2^, resulting in opposing effects between low and high doses of the same agent. For example, synthetic oleanane triterpenoids exert chemopreventive functions at low doses but are also able to induce oxidative stress and apoptosis at higher doses^28, 33^. The same model could be applied to OBT where lower doses of the compound lead to a strong Nrf2 activation while at higher doses, OBT acts as a potent pro-oxidant that targets cancer cells leading to DNA damage and apoptosis as we have previously shown^8^. Given the role of Nrf2 activation in neuroprotection^19, 34, 35^, and the ability of OBT to cross the blood-brain-barrier, low doses of OBT might prove therapeutically beneficial in neurodegenerative disorders for instance, although this remains to be tested.

In conclusion, we have established a mechanistic understanding of the mode of action of OBT. We show that OBT is a reversible covalent modifier of cysteine-residues in Keap1, targeting it for proteosomal degradation, leading to strong activation of Nrf2. In addition, we have developed novel OBT analogs with improved in vivo activity that allow the tuning of pharmacological properties, which will facilitate the preclinical develop of the compound class. Finally, since impaired Keap1 activity and Nrf2 activation leads to increased expression of antioxidant and detoxification genes, OBT and its analogs could potentially have cross-disease applications for disorders such as diabetes, Alzheimer’s disease and Parkinson disease^36^.

## METHODS

### Reagents

OBT was purchased from Gaia Chemicals. N-acetyl-L-cysteine, dithiothreitol, L-glutathione, tert-butylhydroquinone, N-ethylmaleimide, dithiothreitol, iodoacetamide, Cinnamaldehyde, diethyl fumarate and sulforaphane were purchased from Sigma-Aldrich. Trolox, sulfasalazine, CDDO-Me and MG-132 were purchased from Cayman Chemical. Non-targeting shRNA control (shCtrl) and shRNA constructs targeting Keap1 (TRCN0000156676) and CUL3 (TRCN0000012778) were purchased from Sigma and packaged into lentiviral vectors using standard protocols.

### Cell culture

MDA-MB-231 and U87 (U87-MG) cells were obtained from the American Type Culture Collection (ATCC). 293T human embryonic kidney fibroblasts were provided by Dr. Xandra Breakefield (Massachusetts General Hospital). All three cell lines were grown in Dulbecco’s modified Eagle medium supplemented with 10% fetal bovine serum (Gemini Bio-products), 100U penicillin, and 0.1mg/mL streptomycin (Sigma). All cells were maintained at 37°C in a humidified 5% CO_2_ incubator. GSCs were obtained from tumor tissues of GBM patients following surgical resection, under approval from the corresponding Institutional Review Board. These cells have been previously characterized^37–39^ and were maintained in culture as neurospheres in Neurobasal medium (Gibco) supplemented with heparin (2 μg/ mL; Sigma) and recombinant EGF (20 ng/mL) and bFGF-2 (10 ng/mL; Peprotech). Human Astrocytes were obtained from ScienCell and cultured in Astrocytes Medium (ScienCell).

### Cell viability

Cells were plated in 96-well plates and treated with the corresponding compounds. Cell viability was measured by adding 25μl/well of CellTiter-Glo (Promega) followed by 10 minutes incubation and transfer to a white 96-well plate. Bioluminescence was quantified using Synergy HTX multimode reader (Biotek).

### Statistical analysis

GraphPad Prism v6.01 software (LaJolla, CA) was used for statistical analysis of all data. A *p-*value less than 0.05 was considered to be statistically significant. For analysis between multiple groups, a two-tailed Student's *t* test (unpaired), ANOVA, and Tukey’s post-hoc test was performed as indicated. All experiments were performed at least in 3-replicates and repeated 3 independent times.

The mass spectrometry RAW files for the differential alkylation experiments were uploaded to the MassIVE proreomics data repository (massive.ucsd.edu). The accession number is MSV000084638.

(Additional experimental procedures are described in the Supplemental Methods)

## Acknowledgments

We would like to thank Dr. Hiroaki Wakimoto for providing primary GBM cells used in our study. We thank the CCIB DNA Core Facility (for sequencing and oligonucleotides synthesis), MGH Vector Core (for producing the viral vectors (supported by NIH/NINDS P30NS04776) as well as 1S10RR025504 Shared Instrumentation grant for the IVIS imaging system.

## SUPPLEMENTAL FIGURE LEGENDS

**Supplemental Figure 1:**
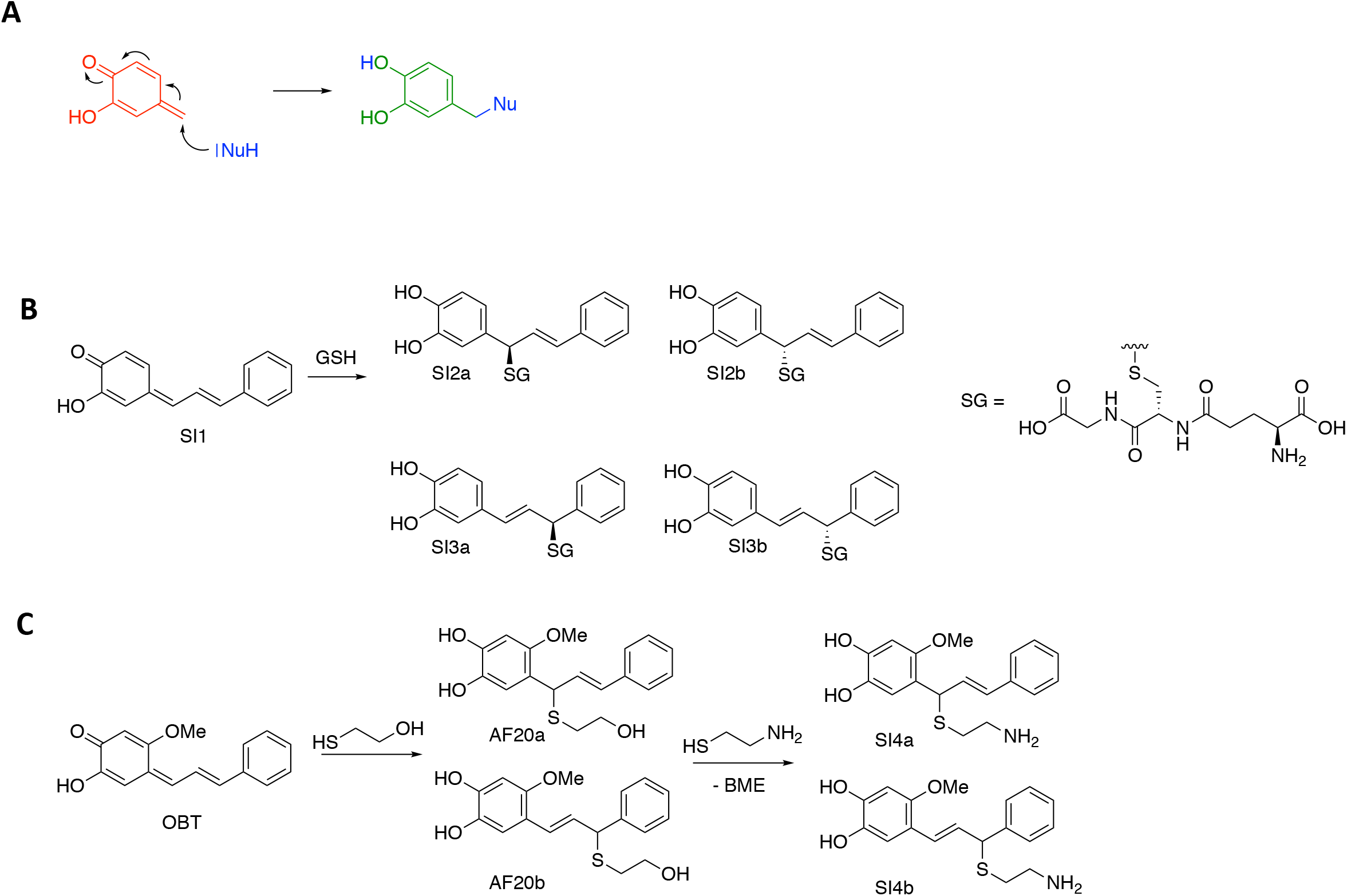
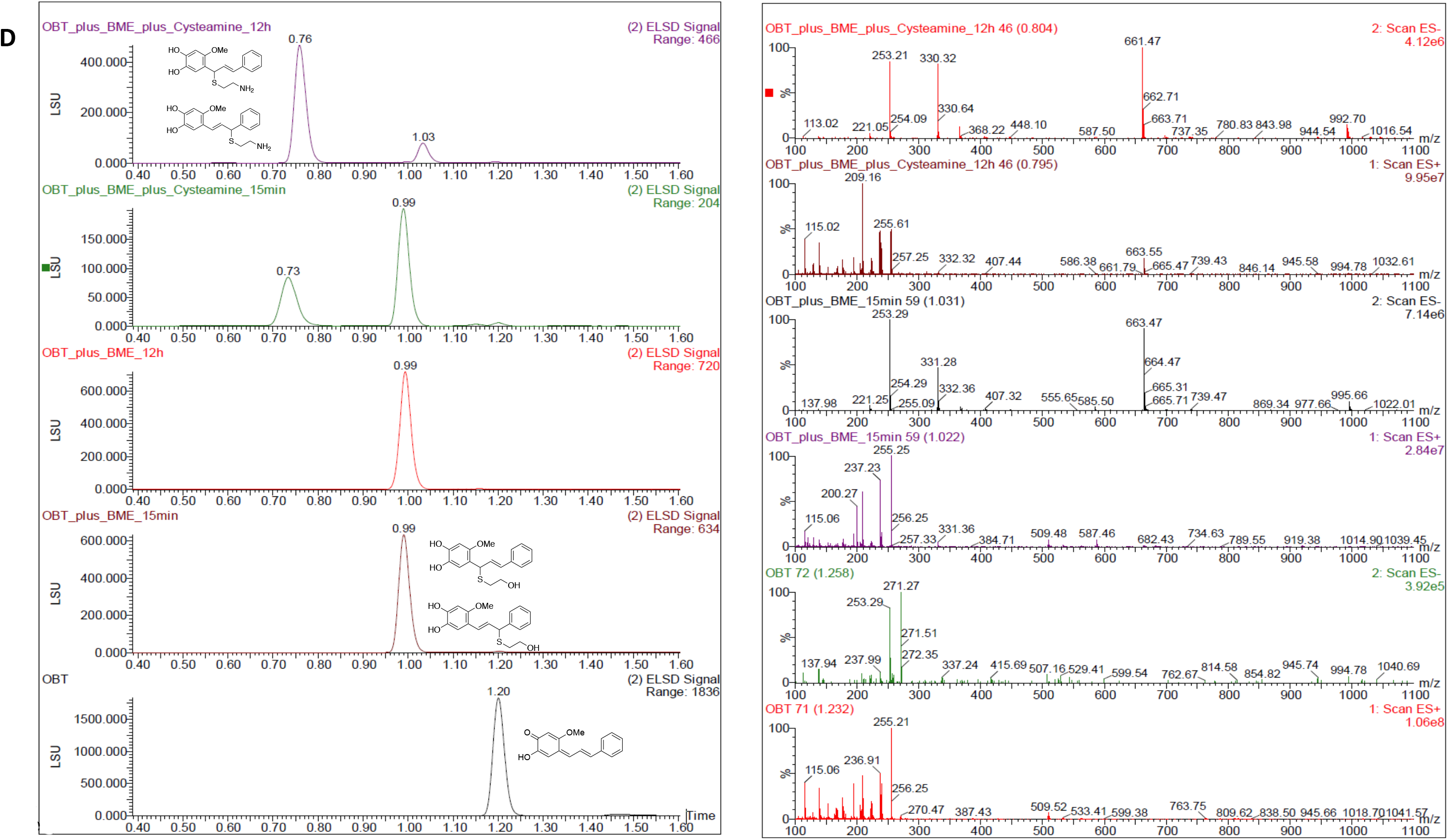
Reactivity of OBT and analogs. **(A)** OBT features a hydroxy para-quinone methide (red), which readily react with nucleophiles (blue) under the formation of catechols. **(B)** Glutathione (GSH) reacts with the quinone methide SI1 under the formation of two distinct diastereomeric pair derived from direct (SI2a, SI2b) and vinylogous (SI3a, SI3b) addition. **(C)** OBT reacts with beta-mercaptoethanol (BME) forming the products derived from direct (AF20a) and vinylogous (AF20b) addition to the quinone-methide core (individual enantiomers are not shown). Addition of excess cystamine to AF20 results in displacement of BME and the formation of SI4a and SI4b. **(D)** LC/MS traces which shows chromatographic separation with baseline separated peaks thus allowing to unambiguously identify and quantify the BME and cystamine adducts of OBT.

**Supplemental Figure 2:**
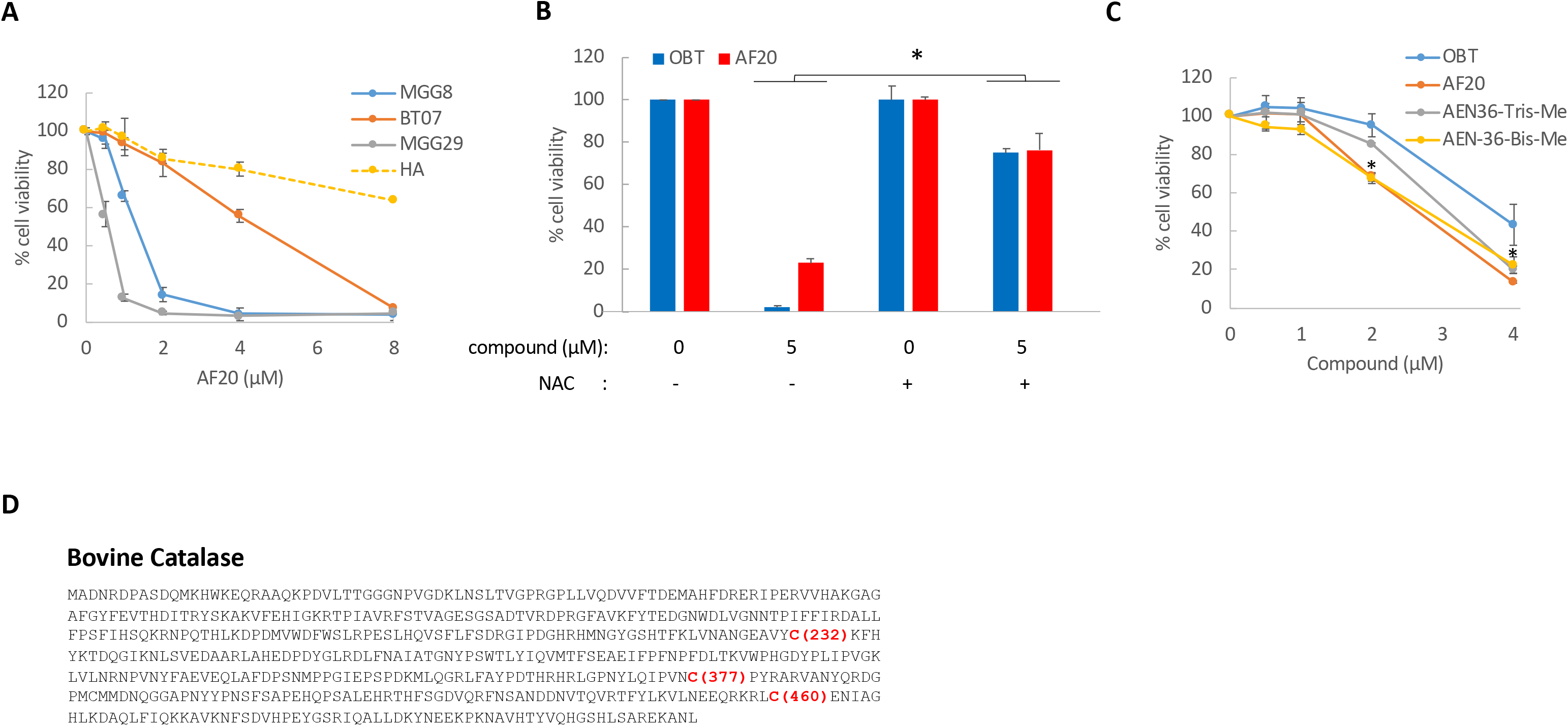
Efficacy of OBT analogs. **(A)** Different GSCs (MGG8; BT07 and MGG29) and normal human astrocytes (HA) were treated with AF20 at the indicated doses. Cell viability was measured four days after treatment and expressed as percentage of control. **(B)** U87 glioma cells were treated with OBT or AF20 in the presence or absence of NAC (3 mM) and cell viability was measured 48h later. **(C)** U87 cells were treated with different doses of OBT or analogs and cell viability was measured four days later and expressed as percentage of vehicle control; *P<0.05 Student *t* test; statistical significance depicts the difference in cell viability between OBT and its analogs. **(D)** Amino acid sequence of the bovine catalase indicating in red the cysteine residue modified by OBT.

**Supplemental Figure 3:**
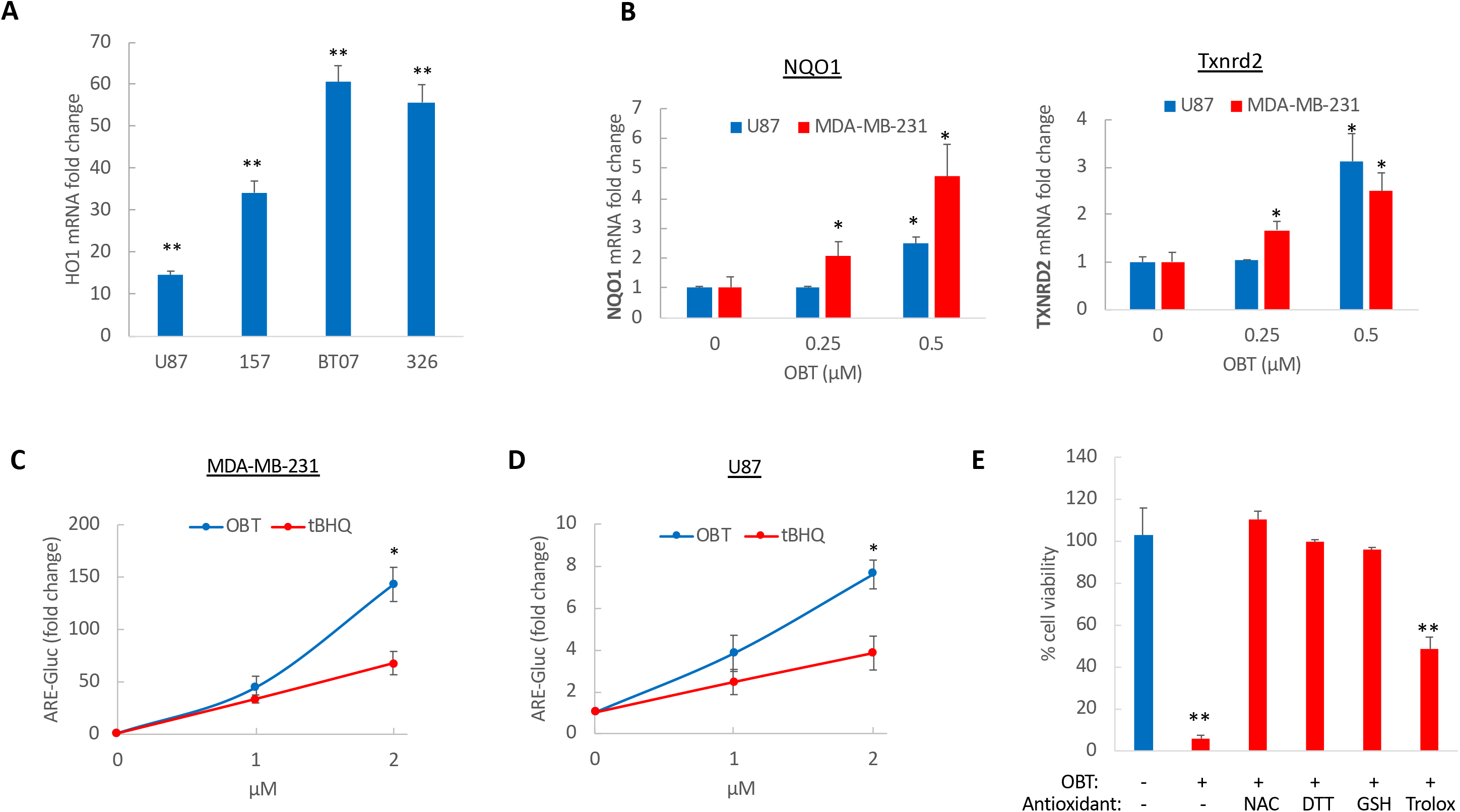
OBT induces Nrf2 activation. **(A)** HO1 mRNA expression levels determined by qRT-PCR and normalized to GAPDH in U87 cells and 3 different GSCs (157; BT12; 326) treated with OBT (1μM) for 8 hours. (**B**) NQO1 and Txnrd2 mRNA levels in U87 and MDA-MB-231 cells after treatment with different doses of OBT. **(C-D)** MDA-MB-231 (C) and U87 cells (D) expressing the ARE-Gluc and SV40-Vluc reporters were treated with OBT or tBHQ. Aliquots of the conditioned medium were assayed for Gluc and Vluc activity. Data presented as a ratio of Gluc/Vluc and normalized to vehicle control (set at 1). **(E)** U87 cells were treated with DMSO (control) or OBT (1μM) and different antioxidants. Cell viability was measured four days later. Data expressed as percentage of cell viability normalized to the control, showing that DTT and GSH completely rescued OBT-mediated cell death, whereas Trolox provided a partial rescue. Statistical significance depicts the difference between the samples treated with OBT alone as compared to the combination of OBT with the antioxidants. *P<0.05; **P<0.001 Student *t* test.

**Supplemental Figure 4.**
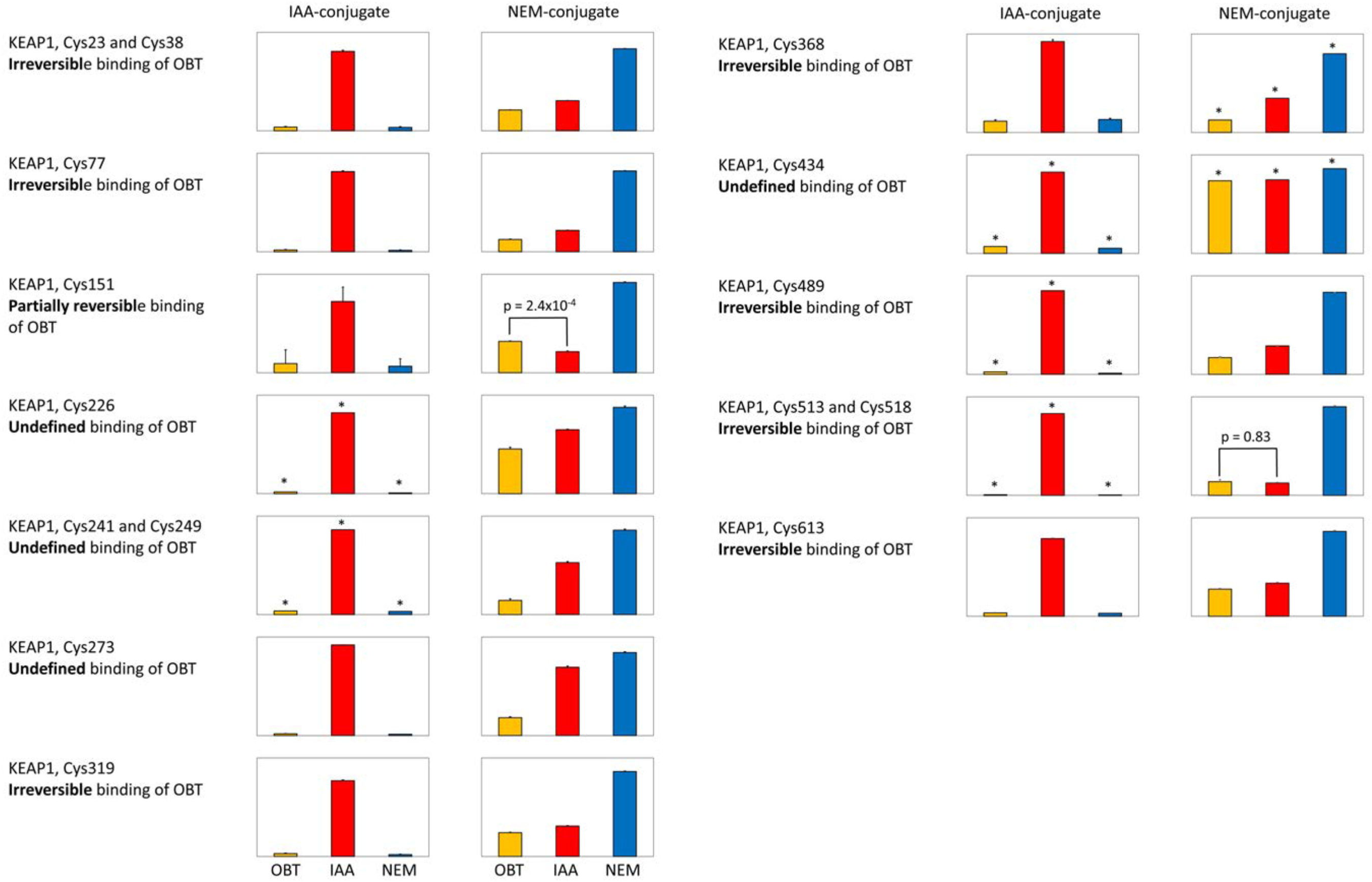
Mapping of OBT covalent biding to cysteine residues in Keap1. Differential alkylation patterns for Keap1 cysteine containing peptides (see Supplemental Table 4). Interpretation guidance of the patterns is given in Figure 1E. The IAA-conjugate pattern defines if OBT is binding to the indicated cysteine-residue; OBT is binding to all twelve quantified cysteines. The NEM-pattern is defining the nature of the OBT binding. We observed three different categories: irreversible binding, with OBT and IAA channels at an intensity at least 2-fold lower than the NEM channel, and the OBT channel not being significantly (p ≤ 0.01, see Supplemental Table 4) higher than the IAA channel intensity; partially reversible binding, as irreversible binding but with an OBT channel intensity significantly higher than the IAA intensity; undefined binding, the IAA channel intensity is at least as high as the NEM channel intensity.

**Supplemental Figure 5:**
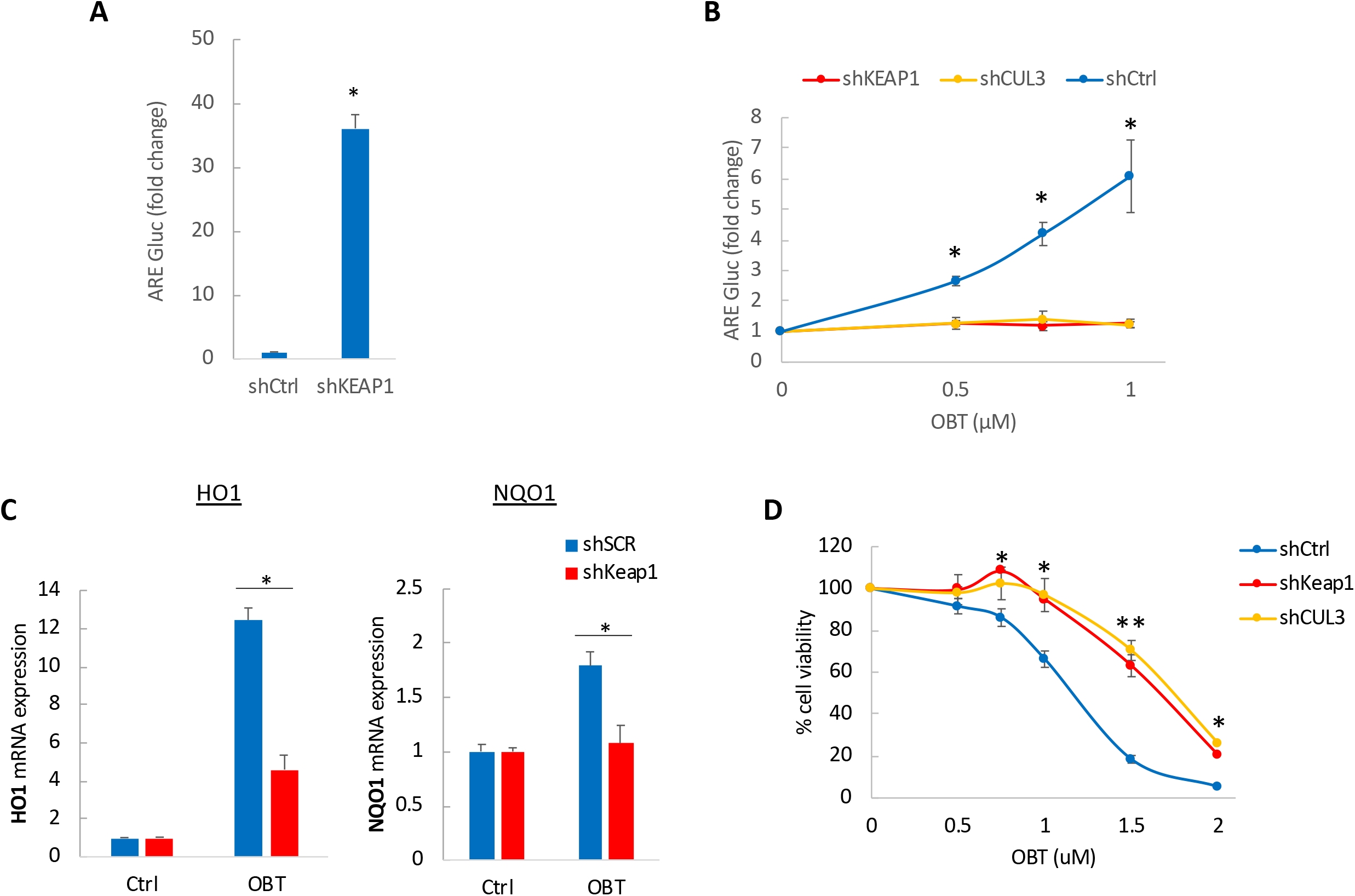
OBT targets Keap1 and activates Nrf2 pathway. **(A)** MDA-MB231 cells expressing ARE-Gluc and SV40-Vluc reporters were transduced with shCtrl or shKeap1. Four days later, aliquots of the conditioned medium were assayed for Gluc and Vluc activity. The data is expressed as the ratio of Gluc/Vluc, normalized to vehicle control (set at 1). **(B)** ARE-Gluc activity normalized to SV40-Vluc in MDA-MB231 expressing shCtrl, shKeap1 or shCUL3 after treatment with different doses of OBT. **(C)** HO1 and NQO1 mRNA expression determined in U87 cells expressing either shCtrl or shKeap1 following 8 hrs of OBT treatment. **(D)** MDA-MB231 cells stably expressing shCtrl, shKeap1 or shCUL3 were treated with different doses of OBT and cell viability was measured after four days. *P<0.05; **P<0.001 Student *t* test.

**Supplemental Figure 6:**
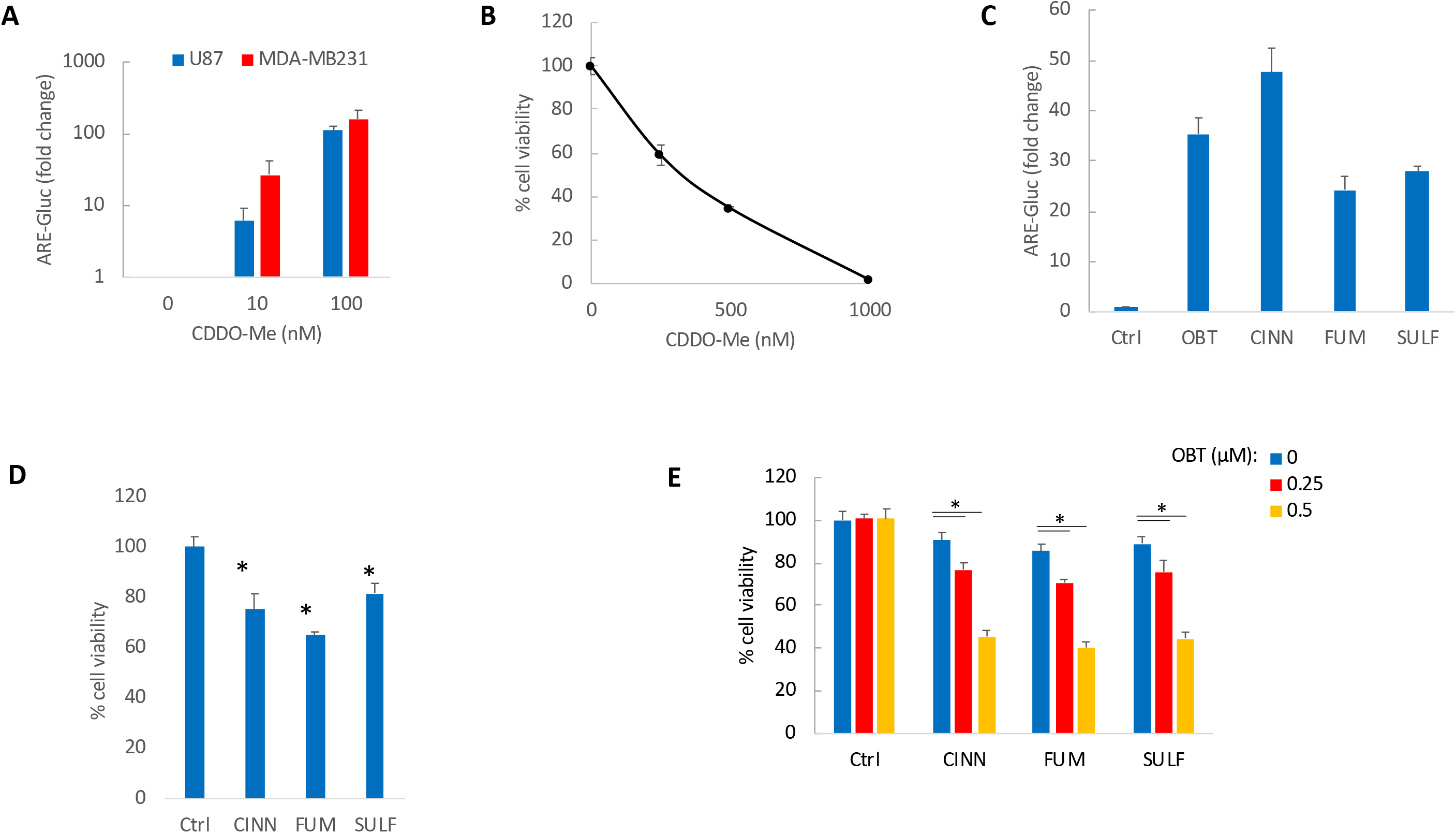
**(A)** U87 and MDA-MB231 cells expressing ARE-Gluc and SV40-Vluc reporters were treated with CDDO-Me at the indicated doses. Aliquots of the conditioned medium were assayed for Gluc and Vluc activity. Data expressed as the ratio of Gluc/Vluc, normalized to vehicle control (set at 1). **(B)** Cell viability of MDA-MB231 cells after 3 days of treatment with CDDO-Me. **(C)** ARE-Gluc activity normalized to SV40-Vluc in U87 cells treated with OBT (1 μM), Cinnamaldehyde (CINN, 100 μM), diethyl fumarate (FUM, 50uM) or sulforaphane (SULF, 5 μM). **(D)** Cell viability in U87 cells treated with CINN, FUM or SULF (same doses as in C). **(E)** U87 cells were co-treated with OBT (at the indicated doses) and either CINN, FUM or SULF (same doses as in C). Cell viability was determined after three days of treatment. *P<0.05 Student t test

**Supplemental Figure 7:**
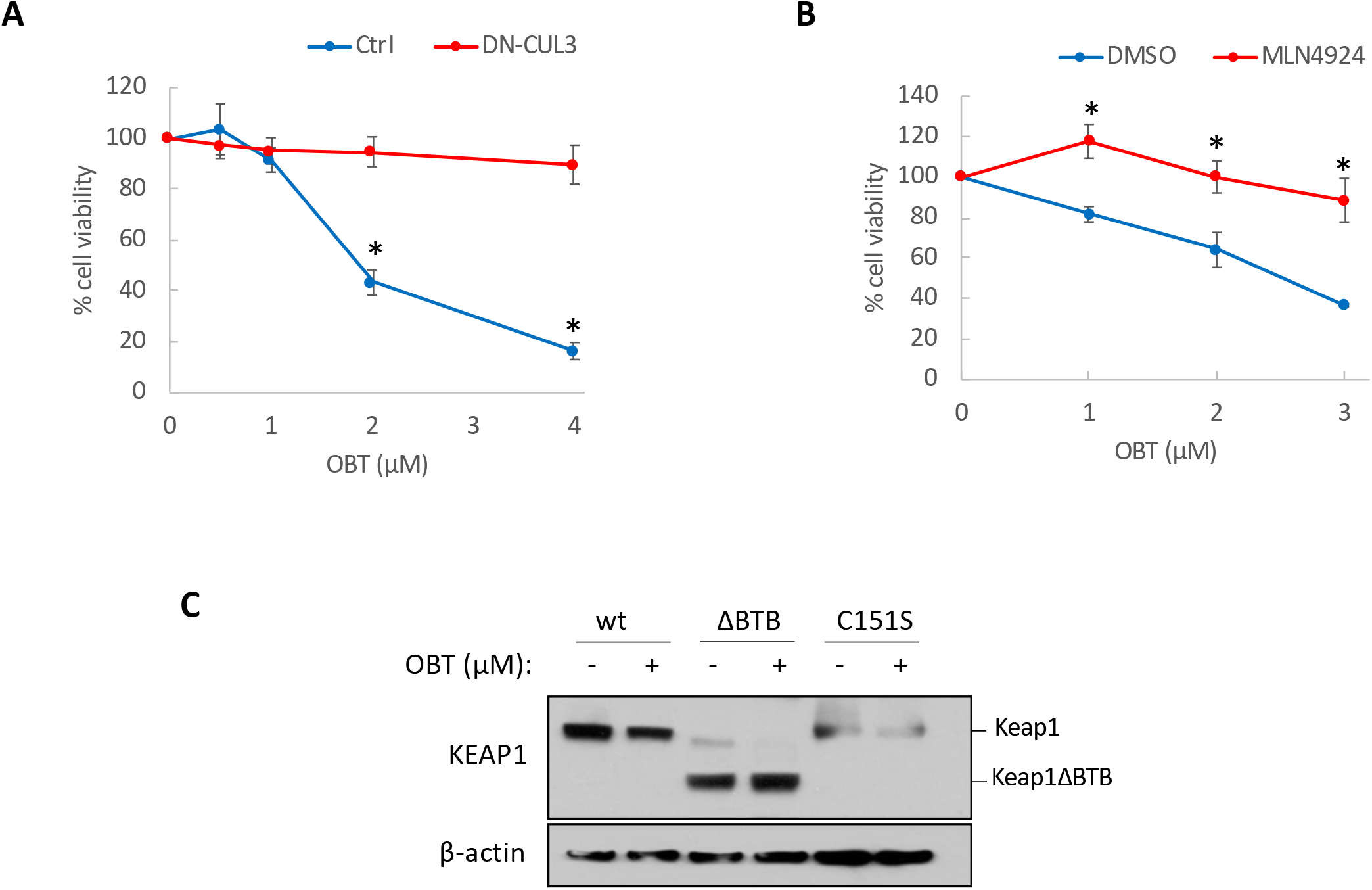
**(A**) MDA-MB231 cells expressing an empty vector (Ctrl) or dominant negative CUL3 (DN-CUL3) were treated with OBT before measuring cell viability after 3 days. **(B)** MDA-MB231 cells were co-treated with OBT and DMSO (control) or MLN4924 (1 μM) and cell viability was measured three days later. **(C)** MDA-MB231 cells were transfected with Keap1 wild-type (wt), Keap1ΔBTB or Keap1C151S, and co-treated with CHX (3 μg/mL) and OBT (4 μM) or vehicle control. Cell lysates were collected after 8 hours and immunoprecipitation was performed using anti-HA antibody followed by immunoblotting for Keap1 and HA. *P<0.05 Student *t* test

**Supplemental Figure 8:**
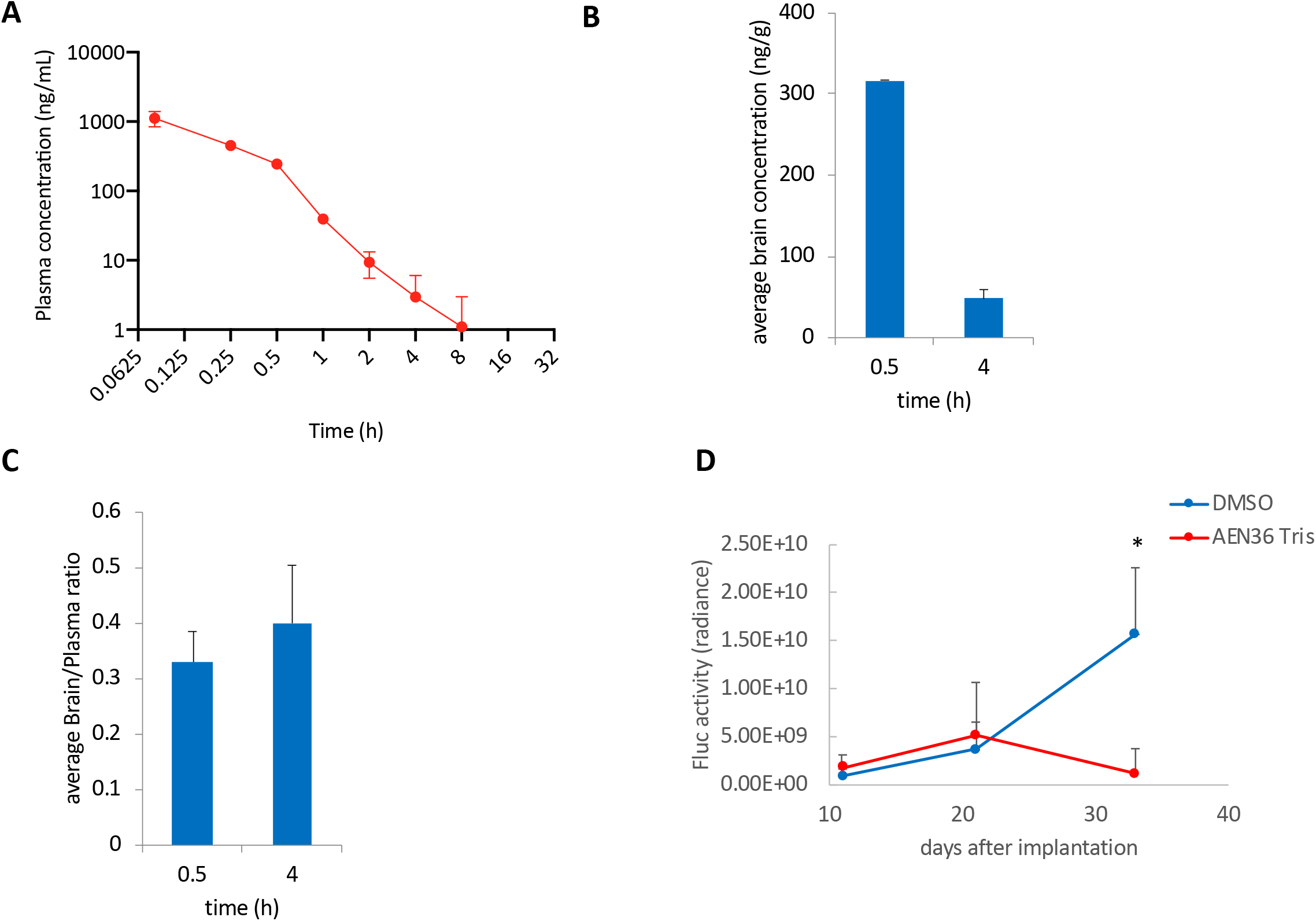
Pharmacokinetics analysis of OBT. **(A)** Average plasma concentration-time profiles of OBT following a single intraperitoneal administration (i.p.; 7.5 mg/kg) in male Swiss Albino mice. Data analyzed by LC-MS/MS. Plasma concentrations were detected up to 24 hrs with Tmax of 0.5 hr. The V*ss* was 3-fold higher than the normal volume of total body water (0.7 L/kg) indicating extravascular distribution. **(B-C)** Average brain concentration (B) and brain/plasma ratio (C) at 0.5 or 4 hours following a single i.p. administration of OBT. The brain-to-plasma ratios ranged from 0.27 to 0.5 at 0.5 hr and 4 hrs respectively. **(D)** Mice-bearing fat pad MDA-MB231 tumors expressing Fluc were treated with either DMSO vehicle control (n=5) or AEN36 Tris (10mg/kg) daily over 22 days. At different time points, tumor volume was monitored by Fluc bioluminescence imaging, and tumor-associated photons were calculated and presented as the average radiance ± SD. *P<0.05 Student *t* test.

## SUPPLEMENTAL TABLE LEGENDS

**Supplemental Table 1:** Bovine catalase peptides quantified in differential alkylation experiment (Fig 1 C-F).

*z*: peptide ion charge.

*XCorr*: primary score of the SEQUEST database search algorithm used to annotate the MS2 spectra.

*dCn*: the secondary score of the SEQUEST algorithm; the relative difference (0-1) between the XCorr of the best match for the MS2 spectra and that of the second best unique peptide sequence.

*Mass deviation*: deviation of the measured intact peptide mass relative to the predicted mass (the deviation is calculated after re-calibrating the acquired data based on high-confidence MS2 annotations);

*LDA_probablility*: The probability of an incorrect peptide annotation calculated using a posterior error histogram from sorting peptide annotations from the forward and the reversed database based on their LDA score.

*Normalized Intensities (OBT1-3, IAA1-3, NEM1-3)*: A two-tiered normalization was performed upon extracting TMT reporter ion intensities form the MS3 spectra. First, an average intensity was calculated for each peptide across all nine channels OBT1-NEM3) and the peptide intensity were normalized using the median average value across all peptides as reference. In a second normalization step, the median intensity for all non-cysteine-containing peptides was calculated for each of the nine channels and used for normalization of all peptides (cysteine-containing and non-cysteine-containing) using the average of all nine median values as reference. Unmarked cysteine residues in the peptide sequences were identified as being alkylated with iodoacetamide (IAA), those marked with a number sign (#) were identified as being alkylated with NEM.

**Supplemental Table 2:** Cysteine containing catalase peptides quantified both with IAA and NEM thiol alkylation in in differential alkylation experiment (Fig 1F). Reporter ions were combined through summation, if peptides including the listed cysteines were quantified more than once in the experiment (see Supplementary Table WH1). Average, standard deviation, and p-values (student’s t-test, two-tailed, unequal variances) were determined across the three measurements performed for each alkylation protocol (OBT, IAA, NEM).

**Supplemental Table 3.** Quantitative mass spectrometry-based proteomics using 10-plexed tandem mass tags (TMT) to simultaneously map protein concentration changes of 7,904 proteins in two GSCs, GBM8 and BT07, either untreated (duplicates) or treated with OBT (triplicates) for 20 hours.

**Supplemental Table 4.** Keap1 peptides quantified in differential alkylation experiment (see Supplemental Table 1 for detailed description of column headers).

**Supplemental Table 5.** Cysteine containing Keap1 peptides quantified both with IAA and NEM thiol alkylation in in differential alkylation experiment (Fig. 2G and Supplemental Fig. 1). One sample was analyzed for each alkylation protocol (OBT, IAA, NEM) and average, standard deviation, and p-values (student’s t-test, two-tailed, unequal variances) were determined all peptides carrying the indicated cysteine residue. If the measurement was done based on only one peptide the average value is the measured intensity and standard deviation

## SUPPLEMENTAL METHODS

### DNA expression constructs

ARE reporter was designed and generated based on our previously published reporter for monitoring of the transcription factor Nuclear Factor Kappa B (NFkB)^1^. NFkB TRE elements were removed and replaced with four copies of the ARE enhancer sequence. This reporter consists of a CSCW lentivirus vector in which we have introduced two copies of the 1.2 kb chicken beta-globin (HS4) insulator elements, a TATA box acting as a minimal promoter upstream of the secreted *Gaussia* luciferase (Gluc). pcDNA3-HA2-Keap1 (Addgene plasmid # 21556) and pcDNA3-HA2-Keap1 delta BTB (Addgene plasmid # 21593) were a gift from Yue Xiong^2^. pHAGE DN Cullin 3 (DN-CUL3) was a gift from Stephen Elledge (Addgene plasmid # 41913). Constructs expressing constitutively active Gluc, Firefly luciferase (Fluc) and *Vargula* luciferase (Vluc) were previously described^1,3,4^. MISSION shRNA Bacterial Glycerol Stocks for Keap1 as well as a non-target shRNA were obtained from Sigma. Transfection of plasmid DNA was performed using Lipofectamine 2000 (Invitrogen) as per manufacturer’s guidelines. Lentivirus packaging and titration was performed in 293T cells by the MGH viral vector core facility following standard protocols. Keap1 mutant C151S plasmid was kindly provided by Dr. Wooyoung Hur.

### Luciferase assay

Coelenterazine (20 μM; Nanolight), The Gluc substrate was added to a 25 μL aliquot of conditioned medium and photon counts were measured using a luminometer. Similarly, the secreted Vluc activity was measured by adding its substrate vargulin (5ng/mL; Nanolight). Cells stably expressing Gluc under the ARE enhancer sequence and Vluc under the constitutively active SV40 minimal promoter were used to determine the ARE reporter activity. The normalized Gluc/Vluc ratio was used as a marker for ARE activation.

### Pharmacokinetics studies

These studies were performed by SAI Lifesciences using Male Swiss Albino mice. OBT was resuspended in a solution formulation of 10% DMA, 10% EtOH, 10% PEG400 in 70% of (2-hydroxypropyl)-β-cyclodextrin (HPβCD). OBT was administered by intraperitoneal (i.p.) injection at 7.5 mg/kg. Blood samples were collected from mice, at 0, 0.08, 0.25, 0.5, 1, 2, 4, 8 and 24 hr. Immediately after blood collection, plasma was harvested by centrifugation and stored until analysis. After collection of blood, brain samples were collected from each mouse at 0.5, 4 and 24 h. Tissue samples were homogenized using ice-cold phosphate buffer saline (pH7.4) and homogenates were stored until analysis. Total homogenate volume was three times the tissue weight. Plasma samples were quantified by LC-MS/MS method (LLOQ = 1.00 ng/mL) for plasma and brain.

### Immunoblotting and Immunoprecipitation analysis

Antibodies against PDI, GAPDH, HA-tag, Ubiquitin, Keap1 and beta-actin (Cell Signaling) as well as secondary Horseradish peroxidase (HRP) conjugated antibodies: sheep anti-mouse IgG-HRP and donkey antirabbit IgG-HRP (Amersham Pharmacia Biotech) were used in this study. For protein expression analysis, cells were lysed in RIPA buffer (150 mM NaCl, 50 mM TRIS, pH 8.0, 1% NP-40, 0.5% deoxycholate, 0.1% SDS) supplemented with 1x protease inhibitors cocktail (Roche). Proteins were quantified using a Bradford protein determination assay (Bio-Rad) followed by electrophoresis in 10% NuPAGE Bis-Tris gels (Life Technologies) and transfer to nitrocellulose membranes (Bio-Rad). Membranes were incubated overnight with antibodies in 1-5% non-fat milk powder in PBS/0.5%TWEEN. Proteins were detected with SuperSignal West Pico Chemiluminescent Substrate (Pierce). For immunoprecipitation assays, cells were lysed in RIPA buffer supplemented with protease inhibitors and 2 mM N-ethylmaleimide. Cell lysates were pre-cleared with protein A agarose beads (Cell Signaling) and incubated with anti-HA antibody with gentle rocking overnight at 4°C followed by incubation with protein A agarose beads and subsequent washing and elution steps. Electrophoresis, transfer and immunoblot analysis was performed as described above.

### Real-time qRT-PCR

Total RNA isolation from cultured cells was performed using RNeasy kit (Qiagen), followed by reverse transcription with OneScript cDNA synthesis Kit (ABM). mRNA expression of different genes was then analyzed by quantitative PCR using PowerUp SYBR Green Master Mix (Applied Biosystems) and performed using a QuantStudio3 real-time PCR system (Applied Biosystems). Primer sequences for HO1, NQO1, TXNRD2 were obtained from the MGH primer bank and were as follows: HO1 Forward, 5’-AAGACTGCGTTCCTGCTCAAC-3’ and reverse 5’-AAAGCCCTACAGCAACTGTCG-3’; NQO1 Forward 5’-GAAGAGCACTGATCGTACTGGC-3’ and reverse 5’-GGATACTGAAAGTTCGCAGGG-3’ and Txnrd2 forward 5’-CTAGCCCCGACACTCAGAAGA-3 and reverse 5’-GGCCATGATCGCTATGGGT. Oligonucleotides were synthesized by the CCIB DNA Core Facility at Massachusetts General Hospital. Expression of human GAPDH was used to mRNA normalization and relative mRNA expression was calculated using the comparative Ct method.

### *In vivo* Tumor Models

Animal experiments performed by our laboratory were approved by the Massachusetts General Hospital Subcommittee on Research Animal Care. Female athymic nude mice (6-8 weeks) were used in all studies with the exception of pharmacokinetics analysis. For brain tumors xenograft model, GBM8 cells (1×10^5^ cells/mouse) stably expressing Fluc were stereotactically implanted into the left striatum of animals. Following implantation, animals were regularly imaged for Fluc to monitor brain tumor growth. To generate the breast cancer model, MDA-MB-231 cells (2.5×10^5^ cells/mouse) expressing Fluc were mixed with Matrigel 1:1 (v/v) (BD Matrigel) and injected into the mammary fat pad of mice. Tris-OBT was dissolved in PBS and administered intraperitoneally. Tumor volumes were measured using a caliper and calculated according to the following formula: volume = (width)^2^ × length/2. For Fluc bioluminescence imaging, Fluc substrate, _D_-luciferin (Gold Biotechnology) (150mg/kg body weight diluted in PBS) was injected i.p. and animals were imaged using an IVIS Spectrum optical imaging system (Caliper Life Sciences) under isofluorane gas anesthesia.

### FDG-PET imaging

Mouse PET/CT scans (n=6, 3 per group) were performed sequentially using a custom-designed mouse bed and PET/CT gantry adapter. Mice were injected intravenously through the tail vein with approximately 600 uCi of fluorodeoxyglucose (F-18-FDG) and imaged on the Inveon small animal imaging system (Siemens Healthcare, Malvern, PA) for positron emission computed tomography and computed tomography (PET-CT) ~45 minutes after injection. Isovue 370 (Bracco Diagnostics, East Princeton, NJ) CT contrast was administered through a tail vein catheter at a rate of 10 ul / min for the duration of the 10 min CT scan for a total volume of 200 ul per mouse, prior to the acquisition of PET images. CT images were acquired over 360 projection with a 500 uA 80 kVp cone beam x-ray tube and a flat panel CMOS detector and reconstructed with a modified Feldkamp cone beam reconstruction algorithm (COBRA) (Exxim Computing Company, Pleasanton, CA) into a 110 micron isotropic voxel matrix. The PET scan was acquired over 30 minutes and reconstructed by filtered back projection with Fourier rebinning and a ramp filter with a Nyquist cutoff frequency of 0.5.

The contrast-enhanced CT images were used to identify the location of the tumor and delineate its margins as regions-of-interest (ROI). The same ROIs are then copied to the matching PET images for quantification of the ^18^F-FDG PET signal as the standard uptake values (SUVs). The SUVs of the tumor are then divided by the SUVs of muscle for each mouse to normalize differences between animals.

### Medicinal Chemistry

Obtusaquinone was purchased from Gaia Chemical Corporation. Absolute ethanol (anhydrous) was purchased from Decon Laboratories. β-mercaptoethanol, acetic anhydride, and pyridine were purchased from Sigma-Aldrich. All chemicals and solvents were used without further purification unless otherwise noted.

Thin layer chromatography was performed with precoated aluminum-backed TLC plates (Silica XG Plates, w/UV254, 200 uM, 20×20 cm) obtained from Sorbtech Sorbent Technologies. Visualization of TLC plates was performed with a UVGL-25 Compact UV Lamp (4 watt, 254/365 nm, 115 V ∼60 Hz/0.16 Amps).

Flash column chromatography was performed on a Biotage Isolera Four Flash Purification System equipped with a 200-400 nm diode array detector using Sorbtech Sorbent Technologies Purity Flash Cartridges (Spherical Silica Gel 12 g, 20-45 uM, 70 A).

Purity of compounds was determined by analytical LC-ELSD-MS performed on a Waters 2545 HPLC equipped with a 2998 diode array detector and a Waters 3100 ESI-MS module, using a XTerraMS C18 5 μm, 4.6 × 50 mm column at a flow rate of 5 mL/min with a linear gradient (95% A / 5% B → 100% B with 90 s and 30 s hold at 100% B, solvent A = water + 0.1% formic acid, solvent B = acetonitrile + 0.1% formic acid).

^1^H and ^13^C NMR spectra were recorded on a Bruker Ascend™ spectrometer at 400 and 100 MHz, respectively. Chemical shifts for protons are reported in parts per million (ppm) and are referenced to residual solvent peaks for CHCl_3_ (7.26 ppm). Data is reported as follows: chemical shift, multiplicity (s = singlet, d = doublet, t = triplet, q = quadruplet, m = multiplet, br = broad), coupling constants (Hz), and integration.

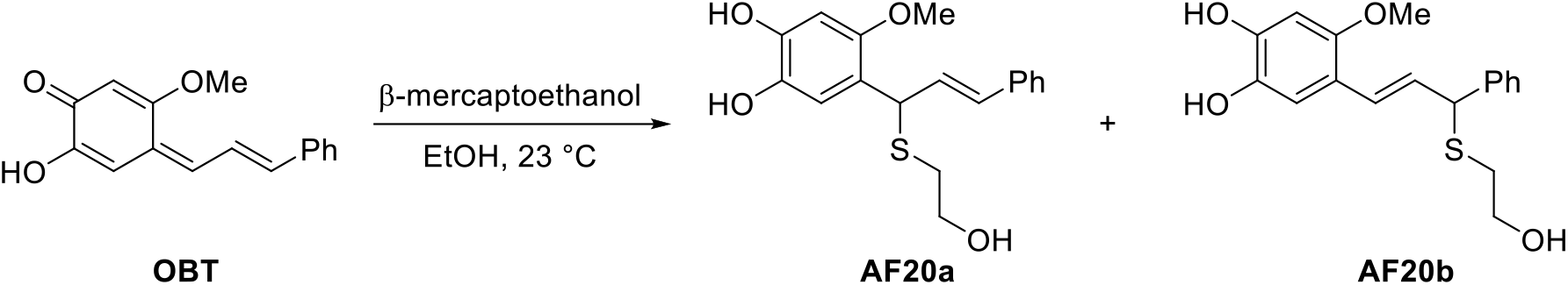

(*E***)-4-(1-((2-hydroxyethyl)thio)-3-phenylallyl)-5-methoxybenzene-1,2-diol** (**AF20a**) and (*E***)-4- (3-((2-hydroxyethyl)thio)-3-phenylprop-1-en-1-yl)-5-methoxybenzene-1,2-diol** (**AF20b**): To a 20-mL flask containing a magnetic stir bar was added obtusaquinone (10 mg, 0.0393 mmol), absolute EtOH (2 mL), and β-mercaptoethanol (8 uL, 0.114 mmol). Over the course of the reaction, the sparingly-soluble orange solid is converted to the adduct, which is homogenous in absolute EtOH and gives rise to a light yellow homoegenous solution. After 1 h, analysis by LCMS indicated that the reaction was not complete, aditional β-mercaptoethanol (4 uL, 0.057 mmol) was added. At t = 2 h, the reaction was complete. The solution is concentrated and dried under high vacuum to yield the desired product as amber oil with AF20a as the major regioisomer (~10:1) Regioisomer A: 1H NMR (400 MHz, MeOD) δ 7.36 (d, J = 8.0 Hz, 2H), 7.26 (t, J = 7.5 Hz, 2H), 7.18 (d, J = 7.4 Hz, 1H), 6.89 (s, 1H), 6.46 (s, 1H), 6.45 (d, J = 15.6, 1H), 6.33 (dd, J = 15.6, 8.4 Hz, 1H), 5.02 (d, J = 8.4 Hz, 1H), 3.74 (s, 3H), 3.62 (t, J = 7.2 Hz, 2H), 2.59 – 2.43 (m, 1H). 13C NMR (101 MHz, MeOD) δ 151.6, 146.3, 140.2, 138.4, 131.5, 131.0, 129.6, 128.5, 127.4, 120.6, 116.3, 101.7, 62.4, 57.0, 45.3, 34.8.

MS (ESI^-^) *m/z* (M-H)^-^ 331.28, [calculated C_18_H_19_O_4_S: 331.1].

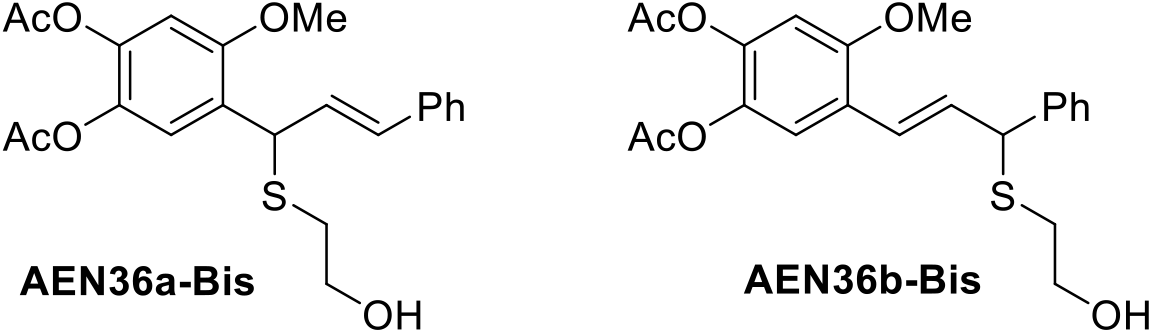

(*E***)-4-(1-((2-hydroxyethyl)thio)-3-phenylallyl)-5-methoxy-1,2-phenylene diacetate** (**AEN36a-Bis**) **and (***E***)-4-(3-((2-hydroxyethyl)thio)-3-phenylprop-1-en-1-yl)-5-methoxy-1,2-phenylene diacetate** (**AEN36b-Bis**): To a 20-mL flask containing a magnetic stir bar and AF20 was added pyridine (786 uL, 0.05 M) and Ac_2_O (7.4 uL, 0.786 mmol). After 16 h, additional Ac_2_O (1.5 uL, 0.0159 mmol) was added. At t = 18.5 h, the reaction was completed as judged by LCMS. The reaction mixture was concentrated to afford an amber oil. Purified by flash column chromatography (0→5% MeOH/CH_2_Cl_2_) to afford a 1:0.24 mixture of **AEN36a-Bis** and **AEN36b-Bis** (5.2 mg, 32%) as a yellow oil, which eventually equilibrated at room temperature in CDCl3 to a 1:1.4 mixture of both isomers. Regioisomer A: ^1^H NMR (400 MHz, CDCl_3_) δ 7.47 – 7.19 (m, 6H), 6.73 (s, 1H), 6.56 (d, *J* = 15.6 Hz, 1H), 6.33 (dd, *J* = 15.7, 8.2 Hz, 1H), 5.07 (d, *J* = 8.4 Hz, 1H), 3.86 (s, 3H), 3.72 (t, *J* = 5.9 Hz, 2H), 2.71 (m, 2H), 2.29 (s, 3H), 2.27 (s, 3H).

^13^C NMR (101 MHz, CDCl_3_) δ 168.6, 168.2, 154.3, 141.5, 136.4, 135.4, 131.7, 128.6, 127.8, 127.6, 127.0, 126.5, 122.9, 106.4, 60.5, 56.2, 43.6, 34.8, 20.7, 20.6.

Regioisomer B: ^1^H NMR (400 MHz, CDCl_3_) δ 7.46 – 7.19 (m, 6H), 6.77 (s, 1H), 6.74 (d, *J* = 15.8 Hz, 1H), 6.31 (dd, *J* = 15.8, 9.1 Hz, 1H), 4.64 (d, *J* = 9.1 Hz, 1H), 3.81 (s, 3H), 3.72 (t, *J* = 5.9 Hz, 2H), 2.82 – 2.58 (m, 2H), 2.29 (s, 3H), 2.27 (s, 3H).

^13^C NMR (101 MHz, CDCl_3_) δ 168.7, 168.2, 154.6, 141.8, 140.1, 135.4, 130.6, 128.8, 127.9, 127.7, 124.7, 124.0, 120.9, 106.3, 60.6, 56.0, 52.2, 34.8, 20.7, 20.6.

MS (ESI^+^) *m/z* (M+Na)^+^ 439.12, [calculated C_22_H_24_NaO_6_S: 439.1].

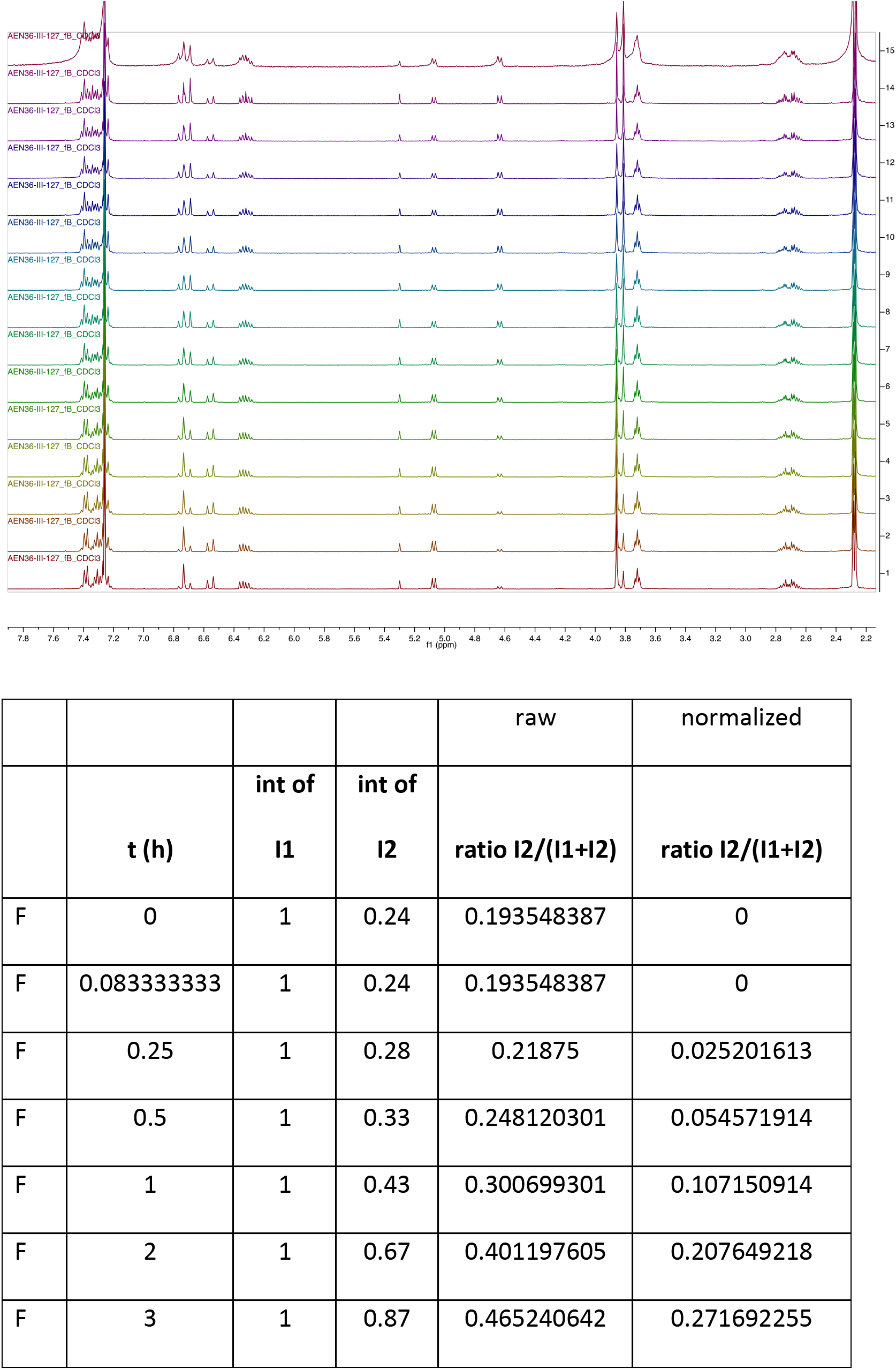
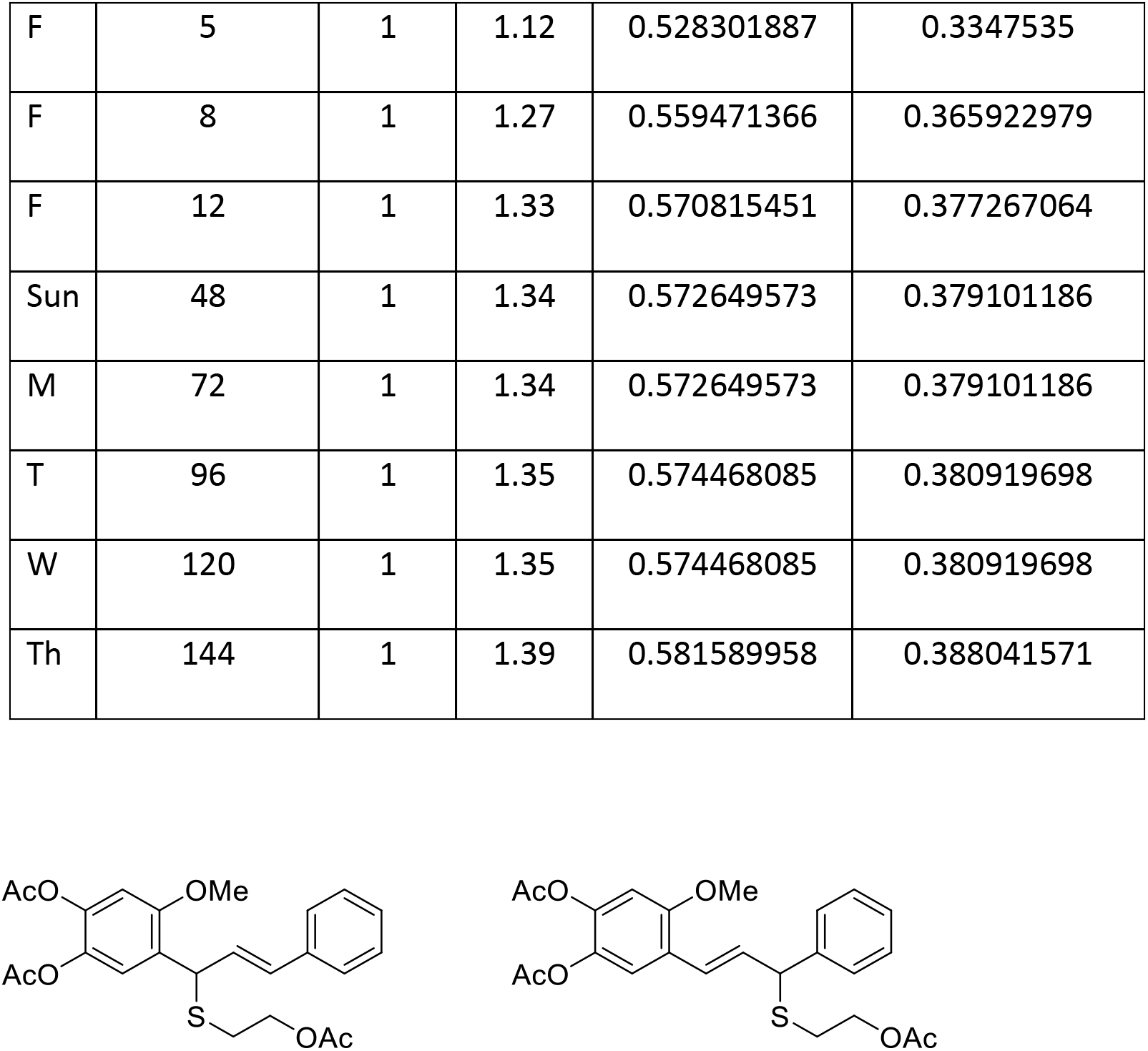

**(E)-4-(1-((2-acetoxyethyl)thio)-3-phenylallyl)-5-methoxy-1,2-phenylene diacetate** (**AEN36a-Tris**) **and (***E***)-4-(3-((2-acetoxyethyl)thio)-3-phenylprop-1-en-1-yl)-5-methoxy-1,2-phenylene diacetate** (**AEN36b-Tris**): To a 20-mL flask containing a magnetic stir bar and 155 mg (0.39 mmol) AF20 was added 5mL pyridine and 160 uL Ac_2_O (1.6 mmol) and stirred overnight. The solution was concentrated and dried under high vacuum to yield the desired product as amber oil with AEN36a-Tris as the major regioisomer (~10:1). Flash column chromatography (0→10% MeOH/CH_2_Cl_2_) afforded the desired products as as a yellow oil as mixtures of regioisomers (170mg, 95%)

Regioisomer A: ^1^H NMR (400 MHz, CDCl3) δ 7.38 - 7.23 (m, 6H), 6.73 (s, 1H), 6.54 (d, J = 15.6 Hz, 1H), 6.29 (dd, J = 15.6, 8.6 Hz, 1H), 5.12 (d, J = 8.6 Hz, 1H), 4.29 – 4.16 (m, 2H), 3.84 (s, 3H), 2.71 (m, 2H), 2.28 (s, 3H), 2.27 (s, 3H), 2.06 (s, 3H).

MS (ESI^+^) *m/z* (M+H)^+^ 459.3, [calculated C_24_H_26_NaO_7_S: 458.14].

### Differential Alkylation for OBT Binding Characterization

10 μg of bovine catalase (Sigma-Aldrich) were suspended in 100 μl of 50 mM HEPES (pH 8.5)/0.5 % SDS and OBT, iodoacetamide (IAA), or N-ethylmaleimide (NEM) was added to a final concentration of 5 mM and the mixtures were incubated for 30 min at RT in the dark before adding IAA to all three reactions to a final concentration of 10 mM in the sample treated with IAA in the first alkylation step, and of 5 mM in the other two samples. Upon incubation at RT for 30 min in the dark dithiothreitol (DTT) was added to all reaction to a final concentration of 20 mM and the reactions were first incubated at RT in the dark for 15 min and then at 56 ºC for 30 min. Then, NEM was added to all reactions to a final concentration of 100 mM followed by incubation at RT in the dark for 30 min. The experiments were done in triplicate. Samples were analyzed by mass spectrometry as described below.

### Quantitative Proteomics

Cell pellets were lysed in a buffer of 75mM NaCl, 50mM HEPES pH 8.5, 10 mM sodium pyrophosphate, 10 mM sodium fluoride, 10 mM β-glycerophosphate, 10 mM sodium orthovanadate, 10 mM PMSF, Roche complete mini EDTA free protease inhibitors (1 tablet per 20 ml), and 3 % SDS. Lysis was achieved by passing the suspension through a 21-gauge needle 20 times. Proteins were prepared for proteomics analysis as described previously ^5^. Proteins from the lysates or the differential alkylation experiments were reduced with DTT; free thiols were alkylated with iodoacetamide; proteins were purified by MeOH/CHCl_3_ precipitation and digested with Lys-C and trypsin, and peptides were labeled with TMT-10plex reagents (Thermo Scientific) ^6^.

Labeled peptide mixtures were pooled, and for the cell line samples, they were fractionated by basic reversed-phase HPLC as described previously ^5^. Twelve fractions were analyzed by multiplexed quantitative proteomics performed on an Orbitrap Fusion mass spectrometer (Thermo Scientifc) using a Simultaneous Precursor Selection (SPS) based MS3 method ^7^. Proteins from the differential alkylation experiments were not fractionated prior analysis by mass spectrometry. MS2 spectra were assigned using a SEQUEST-based ^8^ proteomics analysis platform ^9^. Cysteine residues were allowed to either carry and IAA (mass increment, 57.021464) or an NEM remnant (mass increment, 229.162932). Peptide and protein assignments were filtered to a false discovery rate of < 1 % employing the target-decoy database search strategy ^10^ and using linear discriminant analysis and posterior error histogram sorting ^9^. Peptides with sequences contained in more than one protein sequence from the UniProt database were assigned to the protein with most matching peptides ^9^. We extracted TMT reporter ion intensities as those of the most intense ions within a 0.03 Th window around the predicted reporter ion intensities in the collected MS3 spectra. For the cell line samples only MS3 with an average signal-to-noise value of larger than 40 per reporter ion as well as with an isolation specificity ^11^ of larger than 0.75 were considered for quantification. Only isolation specificity filetring was used when analyzing the data from the differential alkylaton experiments. A two-step normalization of the protein TMT-intensities was performed by first normalizing the protein intensities over all acquired TMT channels for each protein based on the median average protein intensity calculated for all proteins. To correct for slight mixing errors of the peptide mixture from each sample a median of the normalized intensities was calculated from all protein intensities in each TMT channel and the protein intensities were normalized to the median value of these median intensities. For the samples from the differential alkylation experiments the latter normalization was done only considering peptides not carrying a cysteine residue.

## RAW DATA DEPOSITORY

### Bovine catalase

This is an experiment to use differential cysteine alkylation to map the nature of binding of a cysteine-targeting small molecule OBT (for details of the method, please see the Method Section of the manuscript). The alkylating reagents uses besides the studied drug were iodoacetamide (IAA) and N-ethylmaleimide (NEM). As described in the manuscript the experiment includes three routes of serial alkylation experiments termed OBT, IAA, and NEM. Samples were prepared in triplicate and TMT10-plex reagents were used for labeling as follows: OBT replicate 1 (OBT1), 129c; OBT2, 130n; OBT3, 130c; IAA1, 126; IAA2, 127n; IAA3, 127c; NEM1, 128n; NEM2, 128c; NEM3, 129n.

The RAW file for these data is: ol04291_RA_180809_OBT_TMT1.

A differential alkylation experiment was also performed as described above for bovine catalase but using recombinant human KEAP1 as substrate. One set of experiments (OBT, IAA, NEM) was performed. TMT10-plex labeling was performed as follows: OBT, 129n; IAA, 128c; NEM, 128n.

The RAW file for these data is: wh06977_MB_Chris_TMT7_112415

### Modifications

static: 229.162932 (TMT), 57.02146374 (IAA)

variable: 15.9949146221 (oxidation, M), 68.026215 (NEM, in addition to IAA (static))

